# A Toolbox for *Neisseria meningitidis*: Gene Editing, Complementation and Labelling

**DOI:** 10.1101/2024.03.14.585072

**Authors:** Morgane Wuckelt, Audrey Laurent, Clémence Mouville, Julie Meyer, Anne Jamet, Hervé Lecuyer, Xavier Nassif, Emmanuelle Bille, Vladimir Pelicic, Mathieu Coureuil

## Abstract

The efficient transformation of *Neisseria meningitidis* and *Neisseria gonorrhoeae* facilitates the rapid construction of bacterial mutants with insertion of antibiotic resistance cassettes. However, this strategy limits the construction of strains with multiple mutations. Recent advances in markerless strategies for *Neisseria* species have enabled the construction of mutants without antibiotic resistance markers. However, these innovative approaches have potential limitations related to the selection strategy or the possible occurrence of spontaneous mutations in the selection marker genes. In addition, complementation tools or labelling strategies for *N. meningitidis* are also lacking. In this study, we have introduced new tools for markerless mutation, genetic complementation and labelling in *N. meningitidis*, thus improving the research possibilities for understanding and tackling these pathogens.

## INTRODUCTION

*Neisseria meningitidis* and *Neisseria gonorrhoeae* are the two pathogens of the genus *Neisseria.* The first causes meningitis and acute infections of the blood. The second is responsible for sexually transmitted infections and has developed multiple resistance mechanisms against antibiotics. The overlapping of their niches and the ease with which genetic material can be exchanged between *Neisseria* species are possibly driving the emergence of pathogenic strains and resistance to antibiotics (Laver et al., 2015; Brynildsrud et al., 2018; Baerentsen et al., 2022). Today, the main research priorities for these pathogens concern the molecular mechanisms of host-pathogen interaction, the acquisition of genetic material for virulence or resistance to antibiotic and the development of vaccines. These objectives are facilitated by the simplicity of transformation in *N. meningitidis* and *N. gonorrhoeae* and the high efficiency of homologous recombination, which makes it possible to construct bacterial mutants easily and rapidly using current molecular biology methods. Both pathogens are naturally competent for transformation with exogenous DNA thanks to their type IV pili and the competence factors (Facius et al., 1996; Cehovin et al., 2013; Gangel et al., 2014). PCR-based construction of interruption or deletion mutants using antibiotic resistance cassettes has led the community to neglect the development of molecular strategies and tools until recently, although the use of antibiotic resistance cassettes limits the construction of strains with multiple mutations. Strategies for constructing markerless mutants in *Neisseria* species have only been published in the last two years (Jones et al., 2022; Nyongesa et al., 2022; Seow et al., 2023). The work by Nyongesa *et al*. is based on the first selection of streptomycin-resistant strains while the work by Seow *et al*. is based on the co- transformation of a resistance cassette and that of the DNA of interest. These two strategies are effective for obtaining markerless mutants but they require the insertion of an antibiotic resistance gene into the genome. The strategy developed by Jones *et al*. is based on the use of a toxicity cassette carrying the *galK* gene (Ueki et al., 1996). The galactokinase (GalK) phosphorylates 2-deoxy-galactose (2-DOG) into 2-deoxy-galactose-1-phosphate, a toxic intermediate (Alper and Ames, 1975). Jones *et al*. successfully obtained markerless mutants on plates containing 6.4% 2-DOG, a concentration that could be considered a limitation given the price of 2-DOG at retailers. In addition, all three strategies are subject to spontaneous mutation of their selective marker, an event that is likely to generate false-positive mutants and which limits the use of these tools in models of bacteria with a high mutation rate, as in the case of strain Z5463, one of the model organisms of *N. meningitidis* (Achtman et al., 1988). Finally, Geslewitz *et al*. (Geslewitz et al., 2023) proposed very recently to use a CRISPR interference strategy to inhibit the expression of essential genes in *N. gonorrhoeae.* Besides markerless mutation, there are few genetic complementation tools and they have all been developed for *N. gonorrhoeae*. The two main sets of complementation plasmids were developed by the lab of Hank Seifert (namely pGCC4) (Mehr and Seifert, 1998; and unpublished) where they introduced a *lacI* sequence, a *lac* promoter (P*lac*) and an erythromycin or chloramphenicol resistance gene between the two genes *aspC* and *lctP*. This locus appeared to be less attractive for complementation in *N. meningitidis* because of the presence of two small open reading frames (ORF) between *aspC* and *lctP* (*NMV_1885* and *NMV_1887* in the strain NEM8013). More recently, the lab of Joseph Dillard also developed new complementation plasmids based on pGCC4 (pKH37,39,116) and three new plasmids (pMR32,33,68) which introduce either a *lacI*-P*lac* or a tetracycline repressor and the tetracycline-inducible P57opt promoter/operator (P*tet*) between the two genes *iga* and *trpB* (Ramsey et al., 2012).

Bacteria and protein labelling is also poorly developed in *Neisseria* species. GFP tools have been used to detect fluorescent bacteria, but the *gfp* sequence was introduced on a plasmid (Christodoulides et al., 2000; Mairey et al., 2006). This strategy has two limitations, firstly, the use of plasmid in *Neisseria* is not convenient, and secondly, GFP generates free radicals and induces oxidative stress (Ganini et al., 2017). In 2022, Nyongesa *et al*. demonstrated the possible use of mCherry and that of bioluminescence marker in *Neisseria* species (Nyongesa et al., 2022).

Here, we report the development of new tools for mutation, complementation and labelling in *N. meningitidis*. We first adapted the markerless strategy from Jones *et al*. by generating a new cassette containing an antibiotic selection marker and the two counterselection genes *galK* and *pheS\**. The resulting cassette enables rapid and low-cost selection of markerless mutants with no remaining genomic markers, even in strains with a high mutation rate. We then designed complementation plasmids for the use in *N. meningitidis* and *N. gonorrhoeae* with one new genomic locus between genes *NMV_2142* and *NMV_2143*. Finally, we took advantage of the technology based on human O6-alkylguanine-DNA-alkyltransferase (hAGT) (Keppler et al., 2003) to adapt its use to *N. meningitidis*, enabling covalent staining of bacteria or proteins of interest with a wide range of probes.

## RESULTS

### Markerless mutations

Our strategy for obtaining cheap and easy markerless mutations was to design a cassette carrying a kanamycin resistance marker (*aphA3’*) and two counter-selection genes (*pheS\** and *galK*). The cassette, named APG (for *<u>a</u>ph3’ <u>p</u>heS*\* *<u>g</u>alK*), should be minimally affected by point mutations thanks to its two promoters and two counter-selection markers. We assembled the APG cassette from existing constructions. We first benefited from the one obtained by Gurung *et al*. (Gurung et al., 2017) where the *aphA3’* gene is combined into one synthetic promoterless operon with the mutated *pheS\** gene encoding a PheS_A316G_ variant. This variant renders bacteria sensitive to the phenylalanine analogue p-chloro-phenylalanine (Cl-Phe) due to Ala > Gly mutation (Kast, 1994). We then fused this construction to the one proposed by Jones *et al*. (Jones et al., 2022) where the *galK* gene was under the control of an *opaB* promoter. Expressed *galK* phosphorylates 2-DOG and turns it into a non-metabolizable intermediate, conferring sensitivity to 2-DOG. We assessed the sensitivity to Cl-Phe and 2- DOG conferred by the APG cassette inserted into the genome of the two meningococcal strains used in our laboratory: NEM8013 clone 2C4.3 which is a piliated, adherent and encapsulated serogroup C isolate (Nassif et al., 1993) and Z5463 which is a hypermutator encapsulated serogroup A strain isolated from the throat of a patient with meningitis (Achtman et al., 1988) (**Figure 1A, B**). For this purpose, the APG cassette was fused with the 5’ upstream and 3’ downstream genomic sequence of the *pilV* gene, enabling homologous recombination. After transformation and selection for kanamycin resistance, the resulting *pilV*::APG strains were named 2C4.3 APG and Z5463 APG. Wild type (WT) and mutant strains were resuspended in GC Broth (GCB) from an overnight culture on plates with or without kanamycin and serial dilution were spotted on GCB plates with or without Cl-Phe or/and 2-DOG. Cl-Phe at 20 mM and 40 mM appeared to reduce the growth rate of WT meningococci, with a stronger effect for mutants carrying the APG cassette. It should be pointed out that although Cl-Phe reduces the growth rate of *N. meningitidis*, it does not abolish growth or kill the bacteria. On the contrary, 0.5% and 1% 2-DOG had no effect on the growth of WT meningococci, whereas 2-DOG strikingly reduced the growth of mutants. Finally, we chose to combine 1% 2-DOG with 20 mM Cl-Phe. This combination only slightly affected the growth of strain 2C4.3 WT but reduced that of the strain Z5463 WT, and completely suppressed the growth of mutants carrying the APG cassette.

**Figure 1.**
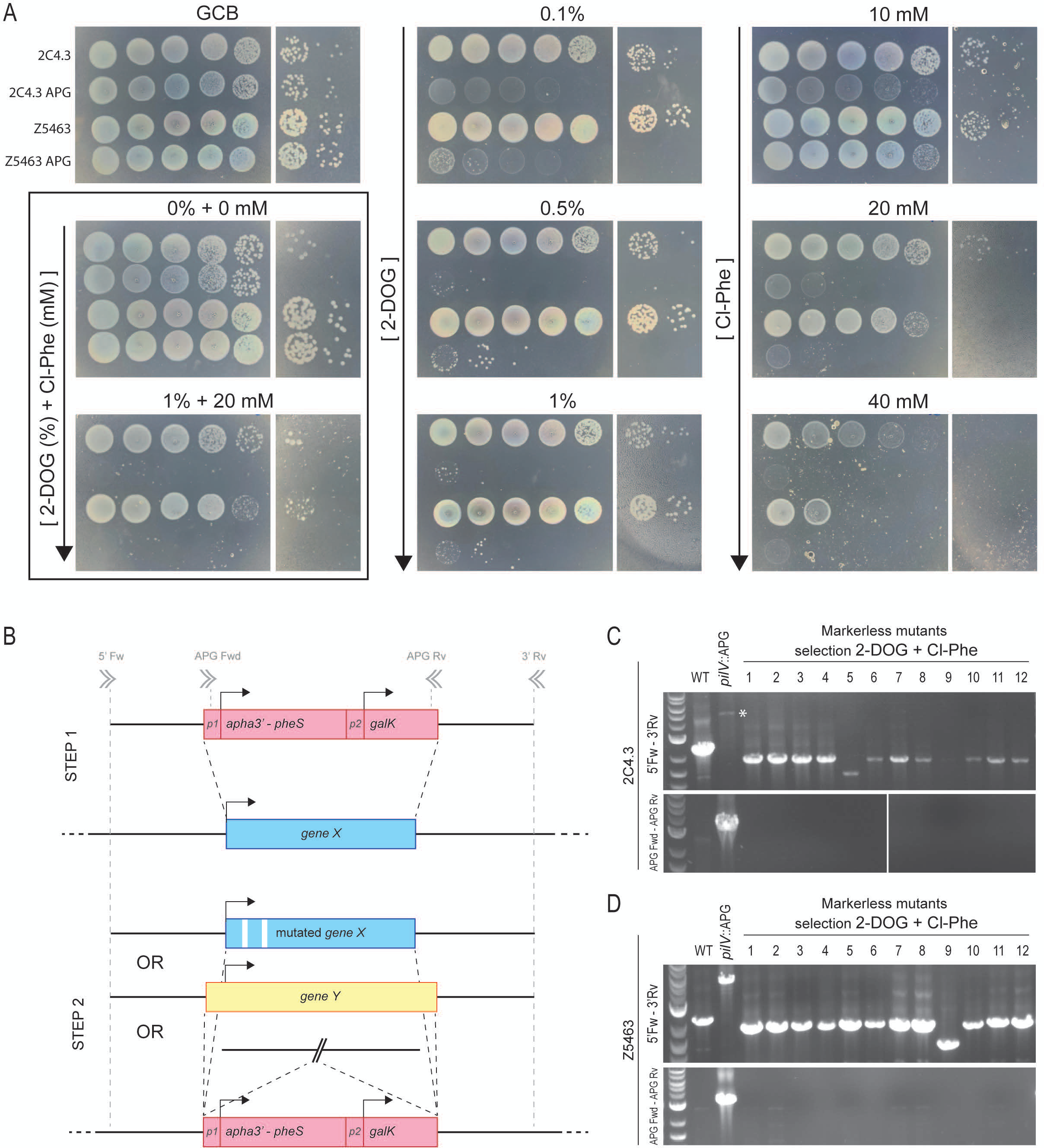
Markerless mutation strategy using 2-deoxy-galactose and p-chloro- phenylalanine. (A) Growth of meningococci on different concentrations of 2-deoxy- galactose (2-DOG) or p-chloro-phenylalanine (Cl-Phe). Wild type strains Z5463 and 2C4.3 and their derivatives carrying the APG cassette were adjusted to 10^9^ CFU/ml and tenfold dilutions were spotted on agar plates supplemented or not with increasing concentrations of 2- DOG or Cl-Phe or both. **(B)** Selection strategy of markerless mutants. In the first step, the genomic DNA sequence of interest is replaced by the APG cassette thanks to homologous recombination. Mutants carrying the APG cassette are selected for the resistance to kanamycin. In the second step, the APG cassette may be removed or replaced by a mutated sequence or another sequence. Markerless mutants are selected on plates containing 2-DOG and Cl-Phe. **(C, D)** Characterization of markerless mutants in the two backgrounds Z5463 and 2C4.3. WT and derivatives carrying the APG cassette in place of the gene *pilV* (*pilV*::APG) and *pilV* markerless mutants obtained by selection on 1% 2-DOG and 20 mM Cl-Phe (2-DOG + Cl-Phe) were checked by PCR using the two primer pairs: 5’Fw-3’Rv and APG Fw-APG Rv. An asterisk indicates the slight band corresponding to the 5’Fw-3’Rv amplicon of the APG cassette in the 2C4.3 background. Clone numbers correspond to those on the plates in Supplementary Figure 1. The largest band on the ladder correspond to 3 kb.

Next, we aimed to remove the APG cassette to obtain a marker-free *pilV* gene deletion mutant. This strategy can also be used to insert a mutated gene or another marker in place of the APG cassette (**Figure 1B**). Strains 2C4.3 APG and Z5463 APG were transformed with the fusion of the 5’ upstream sequence and 3’ downstream sequence of the gene *pilV* to allow homologous recombination. Transformant colonies were picked and isolated on 1% 2-DOG or 20 mM Cl-Phe or a combination of both and grown on GCB agar plates containing either the counter-selective compounds or kanamycin (**Supplementary figure 1**). Transformant colonies that no longer carry the APG cassette should be resistant to 2-DOG and Cl-Phe and sensitive to kanamycin. In any case, Cl-Phe alone was not enough to select a population of markerless mutants. Surprisingly, we observed resistance to both Cl-Phe and kanamycin, suggesting that a compensatory mutation may occur in the *pheS\** gene or elsewhere in the genome, or that the colonies selected were a mix of a strain carrying the APG cassette and a markerless mutant strain. However, a second round of selection by picking and isolating a colony from each of the twelve clones did not allow us to recover kanamycin-sensitive clones with Z5463 strain. The use of 2-DOG alone resulted in the selection of 8 out of 12 markerless populations in the 2C4.3 APG strain, although we observed few kanamycin resistant contaminating colonies. However, for the strain Z5463 APG, the use of 2-DOG alone was not effective for the selection of markerless mutant. A second round of selection was performed and only 2 markerless mutants out of 12 clones were isolated. Besides, the combined use of 2- DOG and Cl-Phe has proved successful. 12 markerless clones out of 12 were obtained with the strain 2C4.3 APG after the first isolation with very few contaminations from kanamycin resistant bacteria. 11 clones out of 12 from the strain Z5463 APG were sensitive to kanamycin with few contaminations from kanamycin resistant bacteria. After a second step of isolation, most of the Z5463 clones were fully sensitive to kanamycin. Genomic DNA were obtained from the 2C4.3 Δ*pilV* markerless (ML) clones and Z5463Δ*pilV* ML clones selected on 2-DOG and 2-DOG/Cl-Phe. Then, markerless mutations were confirmed by PCR (**Figure 1C** and **Supplementary figure 1**) and the growth of one selected mutant was compared to the WT and *pilV*::APG derivative strains (**Supplementary figure 1B**, **C**). In conclusion, while Cl-Phe is not sufficient for the selection of markerless mutants in our hands, its combination with 2- DOG proved to be successful and can be considered for the routine selection of markerless mutant.

### Complementation and expression vectors for N. meningitidis

Once a markerless mutant has been designed, the most appropriate confirmation for the phenotype observed with the mutant strain might be on-site complementation by re-inserting the gene of interest in place of the markerless mutation. However, the use of an inducible promoter may be beneficial in certain circumstances and it may also be convenient to overexpress a gene in a wild-type context. To date, two loci were proposed as genomic insertion sites for *N. gonorrhoeae* and used in *N. meningitidis*: (i) between *lctP* and *aspC* or (ii) between *iga* and *trpB*. The first locus should be avoided in *N. meningitidis* because of the presence of two ORFs between the genes. The second locus is appropriate to use in *N. meningitidis*. However, since *iga* is variable (**supplementary dataset_1**), we decided to clone the 3’ ends of *iga* and *trpB* sequences from the meningococcal strain NEM8013 and then two new plasmids were designed based on pMR33 and pMR68 from Ramsey *et al*. (Ramsey et al., 2012). The two plasmids were named pNM99_Ptet_ and pNM99_Plac_ (**Figure 2A**, **B**). They carry the anhydrotetracycline-inducible promoter (P*tet*) and *tet*-repressor or the lac promoter (P*lac*) and operator and the *lac* repressor, respectively. In order to maintain the possibility of performing double complementation/expression, we then selected a second locus which replaced the *lctP*/*aspC* locus. We have chosen an insertion site between the two genes *NMV_2142* and *NMV_2143* in the NEM8013 strain. These two genes are conserved in *N. meningitidis* and *N. gonorrhoeae* (**supplementary dataset_1**) and are encoded in opposite directions with the two 3’ ends facing each other. Two terminators with two DNA uptake sequences (DUS) each are present between the two genes. As for pNM99_Ptet_ (used as a template), the P*tet* and *tet*-repressor were added between the two genes *NMV_2142/NMV_2143*. The resulting plasmid was named pNM42_Ptet_ (**Figure 2C**). Then, we exchanged the *ermC* genes encoding resistance to erythromycin for the *aadA* genes encoding resistance to spectinomycin in both pNM99_Ptet_ and pNM42_Ptet_ plasmids. Therefore, the combined use of pNM99_Ptet/Plac_ and pNM42_Ptet_ is possible for the same strain.

**Figure 2.**
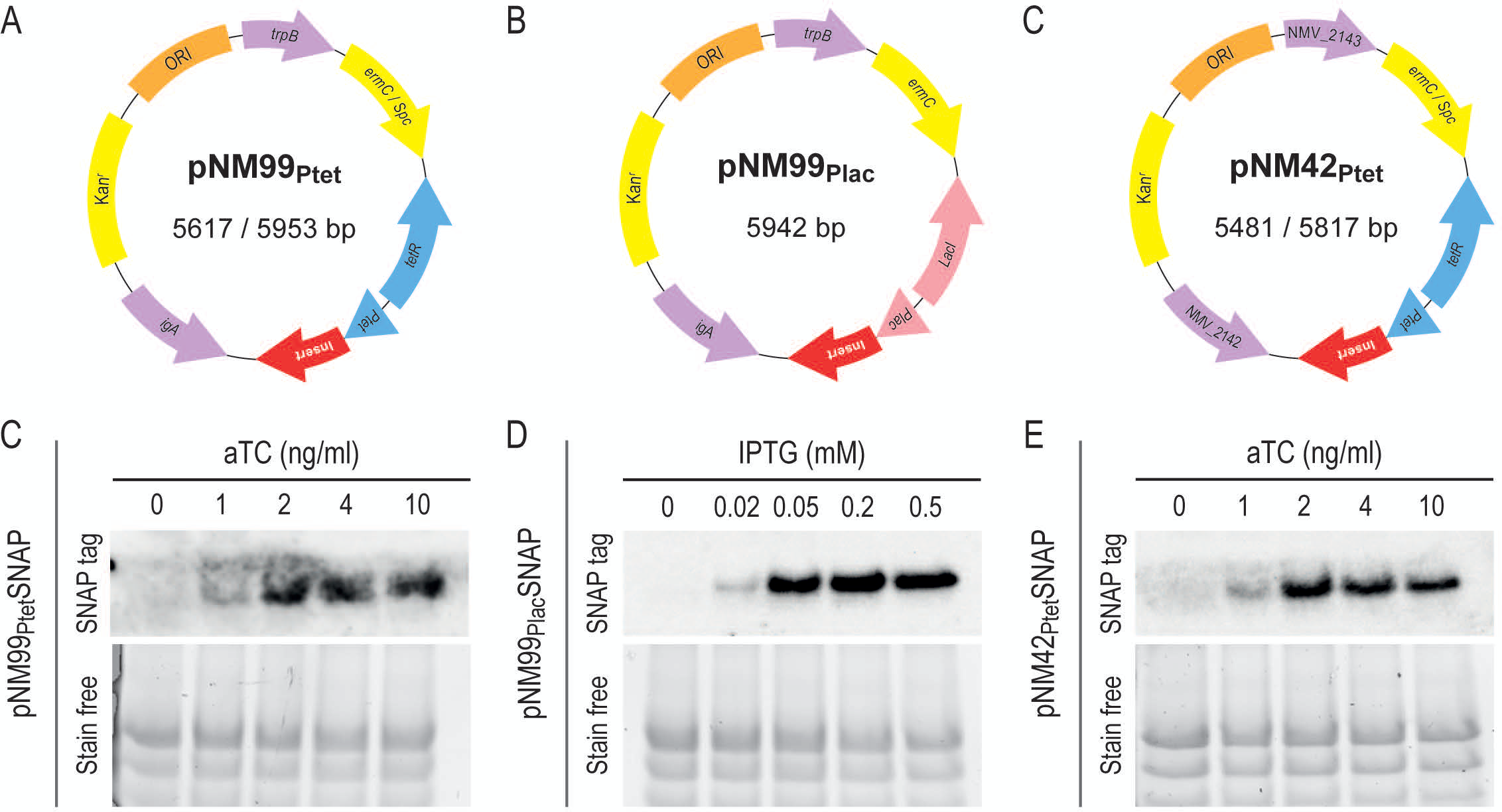
Complementation plasmids. (A, B, C) Circular maps of (A) pNM99Ptet, (B) pNM99Plac and (C) pNM42Ptet. Regions of homology with *N. meningitidis* chromosomes are shown in purple. Antibiotic selection cassettes are shown in yellow. P*lac*: *lac* promoter and operator. LacI: the lac repressor. P*tet*: anhydrotetracycline-inducible promoter. TetR: *tet* repressor. Insertion site is indicated in red. **(C, D, E)** Expression of the SNAP tag using the different plasmids. Expression of the SNAP tag under the control of P*tet* or P*lac* was assessed in WT 2C4.3 transformed with (C) pNM99-_Ptet_SNAP or (D) -_Plac_SNAP or (E) pNM42_Ptet_SNAP. Bacteria were incubated overnight on GCB agar plates containing increasing concentrations of aTC or IPTG. Expression of the tag was determined by western blotting using anti-SNAP tag antibodies. Total proteins were detected in the gel by Stain-Free technology.

Finally, to assess the expression of protein using these constructions, we inserted the *SNAP-tag* sequence obtained from the pSNAP-tag (T7)-2 vector (New England Biolabs) after P*tet* or P*lac*. This sequence encodes a 20 kDa mutant of the DNA repair protein O6-alkylguanine-DNA alkyltransferase that reacts with benzylguanine derivatives, resulting in irreversible covalent labelling of the tag with the dedicated synthetic probe (Keppler et al., 2003). This tag is particularly interesting as it allows the same tagged protein to be labelled with different fluorophores or biotin. In addition, when expressed alone, this tag can be used to stain bacteria. Therefore, 2C4.3 WT bacteria were transformed with pNM99-_Ptet_SNAP or -_Plac_SNAP or pNM42_Ptet_SNAP. Erythromycin-resistant clones that had performed recombination at the locus *iga/trpB* or *NMV_2142/ NMV_2143* were selected and amplified. Bacteria were plated on GCB agar plates supplemented with increasing concentrations of anhydrotetracycline (aTC) or isopropyl β-D-1-thiogalactopyranoside (IPTG), lysed and processed for SNAP tag identification by western blotting. A strong SNAP tag expression was observed with 2 ng/ml aTC and 0.05 mM IPTG (**Figure 2**). These results demonstrate that the two complementation plasmids are functional in *N. meningitidis* and allow strong expression of proteins under the control of P*tet* or P*lac*.

### Covalent staining of bacteria using the hAGT-based technology

We then asked whether the SNAP tag could be used in *N. meningitidis* to stain live or fixed bacteria. To easily manipulate and observe bacteria, we decided to work on bacteria that form colonies on human cells. WT bacteria or bacteria expressing the SNAP tag were grown in the presence of aTC. We infected endothelial cells grown on glass coverslips with the above-mentioned bacteria for two hours and fixed them with 4% paraformaldehyde (PFA). Co-staining was performed in a final volume of 30 µl using Dapi and 0.12 μM of SNAP-Cell TMR-Star, a red fluorescent membrane-permeable substrate of the SNAP tag (**Figure 3A**). This substrate concentration was sufficient to clearly reveal bacteria expressing the SNAP label, whereas it generated a slight background, barely revealing endothelial cells and none of the WT bacteria.

**Figure 3.**
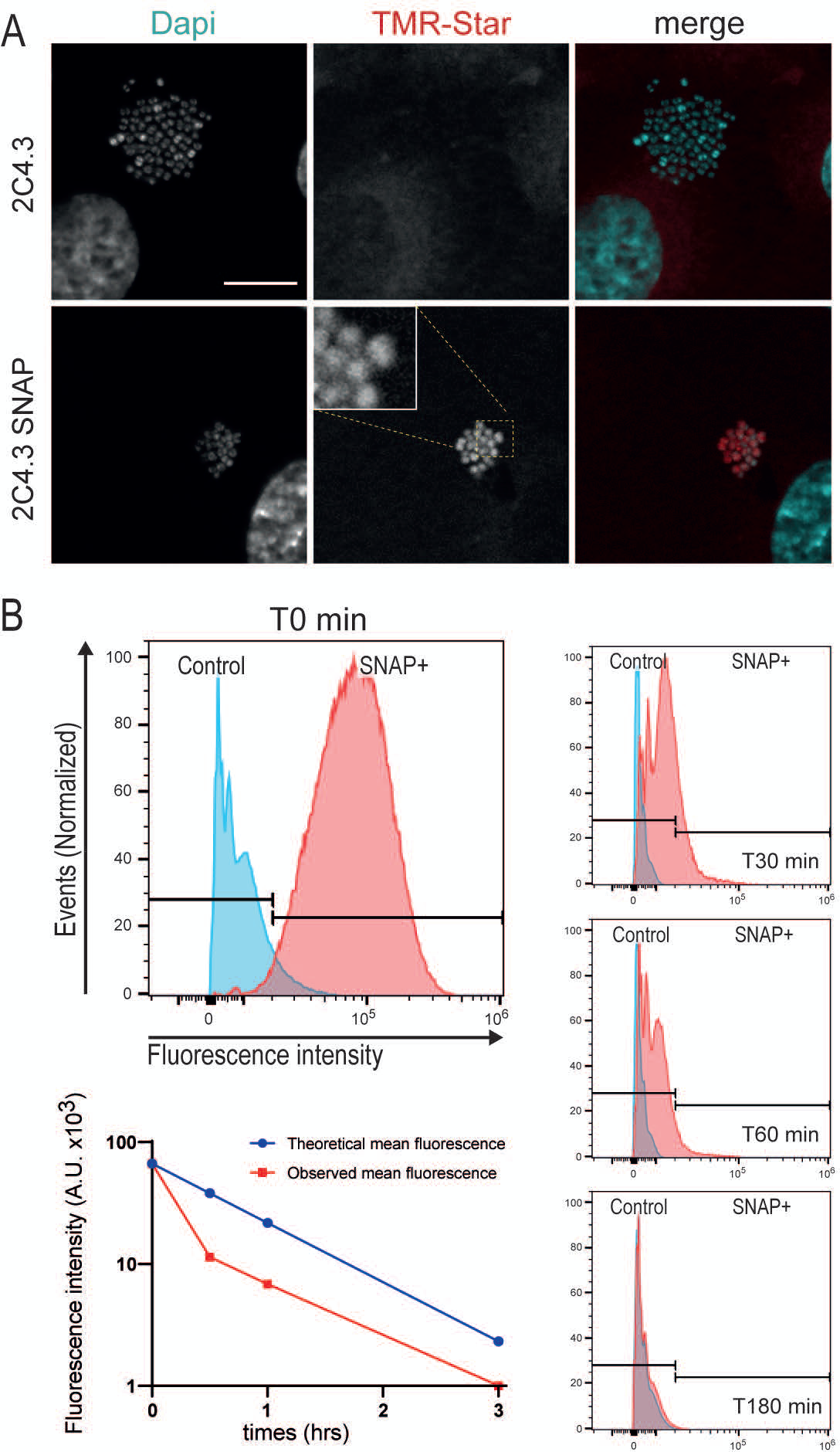
Staining of living and fixed bacteria using the hAGT-based technology. (A) Imaging of bacteria stained with the red fluorescent membrane-permeable substrate TMR-Star after fixation. Endothelial cells were infected with 2C4.3 WT or derivatives expressing the SNAP tag (SNAP), then fixed and processed for staining with Dapi or 0.12 µM of SNAP-Cell TMR-Star. Bar: 10 µm. **(B)** Flow cytometry analysis of TMR-Star fluorescence in bacteria stained before fixation. Bacteria grew in GCB liquid media and were incubated 30 min with the substrate, washed and fixed in 4% PFA (T0 min) or grew again in fresh broth and fixed after 30 min (T30 min), 60 min (T60 min), or 180 min (T180 min). To visualise the decrease in fluorescence, the theoretical mean (blue) fluorescence and the observed mean (red) fluorescence were plotted against time

Next, we asked whether this tag could be used as a marker to follow the growth of living bacteria in a fluorescence dilution assay (Roostalu et al., 2008). In theory, living bacteria stained with the substrate should have their fluorescence reduced by 2 fold per division, which corresponds to approximately 40 minutes (Tobiason and Seifert, 2010). Bacteria were grown in culture media, incubated with the TMR-Star substrate for 30 min and washed. Then, bacteria were either fixed with 4% PFA or grown again for 30 min, 60 min or 180 min before fixation. The fluorescence of bacteria was revealed by flow cytometry. While bacteria that were fixed just after staining could be easily distinguished, the fluorescence decreased very fast with a drop of approximately 90% in signal intensity as soon as 30 minutes after staining (**Figure 3B**). The decrease in fluorescence did not follow the theoretical decrease in fluorescence caused by bacterial division. Therefore, this staining strategy could not be used to monitor the growth of *N. meningitidis*, although it does allow the staining of fixed or living bacteria.

## DISCUSSION

*Neisseriaceae* in general and pathogenic *Neisseriaceae* in particular represent an important area of research, particularly given the severity of diseases caused by meningococcus or the antibiotic resistance of gonococci. They are naturally transformable and easy to handle, although the level of biosecurity must be taken into account. However, few tools applicable to *Neisseriaceae* have been developed for research purposes. In this work, we have adapted three new tools for the study of *N. meningitidis* to (i) perform markerless mutations at reduced cost; (ii) conditionally express one or two genes; and (iii) demonstrate the feasibility of using hAGT-based technology. The close relationship between *Neisseriaceae*, and in particular between the two pathogenic members of this family, suggests that these tools should easily be transferred to the gonococcus or other *Neisseriaceae*.

Our markerless strategy was based on the use of two sensitivity genes. The first, *galK*, sensitises bacteria to 2-DOG, a strategy proposed by Jones *et al*. (Jones et al., 2022) in gonococci. The second, the mutated *pheS*\* gene, sensitises bacteria to Cl-Phe, which had not been tested in *Neisseriaceae* (Gurung et al., 2017). It should be noted that the meningococcus is slightly sensitive to Cl-Phe. In our hands, the mutated gene *pheS\** associated with Cl-Phe is not sufficient to allow selection of markerless mutants. Our observations suggested that we were selecting colonies in which the bacteria were resistant to Cl-Phe but still carried the APG cassette, whether we used strains Z5463 or 2C4.3. In addition, 2-DOG was not toxic to the meningococcus. In strain 2C4.3 expressing *galK*, the use of 2-DOG alone enabled the selection of markerless mutants with a good success rate of 66%, although the mutants were not perfectly pure after single isolation. On the other hand, in strain Z5463, the use of 2-DOG alone failed to select markerless mutant after a single isolation and was poorly effective after two isolation steps.

The appearance of a population of bacteria resistant to kanamycin and losing sensitivity to Cl-Phe or 2-DOG suggests that the population remained chimeric after transformation, consisting of bacteria carrying either the APG cassette or the markerless mutation. Furthermore, we cannot rule out the possibility of compensatory mutations in the *galK* and *pheS\** genes or in other genes. As pathogenic *Neisseria* are able to maintain multiple chromosomes (Tobiason and Seifert, 2010), the APG cassette may be difficult to eliminate following homologous recombination. Furthermore, the concentrations of 2-DOG or Cl-Phe used in our work may not be sufficient to ensure complete elimination. As a result, the bacterial population remains resistant to both kanamycin and 2-DOG or Cl-Phe even after a second isolation.

The combined use of 2-DOG and Cl-Phe proved to be very effective with a high success rate in strain 2C4.3. A second isolation was necessary for the strain Z5463, with a very good success rate. The mutants obtained growth comparable to that of the wild type or the strain carrying the APG cassette. Finally, the use of a cassette combining two susceptibility genes allows easy and efficient selection in *N. meningitidis*, while reducing the concentration of 2- DOG from 6% (Jones et al., 2022) to 1%, thus reducing costs.

The plasmids used to express recombinant genes in pathogenic *Neisseria* were all developed for use in gonococci. Their use in meningococcus has seemed so logical that no plasmid has been developed specifically for this bacterium. However, the use of the *aspC*/*lctP* locus is problematic because of the presence of two ORFs between these two genes in *N. meningitidis*. Here, we have followed the strategy developed by Ramsey *et al*. (Ramsey et al., 2012) to construct a plasmid dedicated to meningococcus, using the sequences of the *igA* and *trpB* genes of strain 2C4.3. In parallel, we proposed a new locus to host a conditional gene expression module, between *NMV_2142* and *NMV_2143*. This locus has the advantage of being highly conserved in both *N. meningitidis* and *N. gonorrhoeae* (**Supplementary dataset_1**).

We used these constructs to evaluate hAGT technology for labelling bacteria or proteins of interest in *N. meningitidis*. The SNAP tag proposed by New England Biolabs is available in two versions (SNAP and CLIP) allowing for dual labelling. We have shown that the SNAP tag is functional for labelling under *in vivo* conditions or after fixation, allowing observation by microscopy or flow cytometry. However, our results indicate that *in vivo* staining requires rapid fixation, as staining intensity fades rapidly in living bacteria. Overall, this proof of concept paves the way for molecular staining strategies for pathogenic *Neisseria*. One of the main advantages of the SNAP tag is its flexibility. The tag can be used for both visualisation and purification from the same pre-prepared sample. This tag allows the use of a marker with a 1:1 stoichiometry between target and marker. The staining is performed at the end of the experiment and not throughout the life of the bacteria, as is the case with GFP. This should limit the effects of oxidative stress on the bacteria. The SNAP tag is therefore compatible with high-resolution microscopy strategies such as dSTORM or STED, as well as electron microscopy and gold bead labelling, thanks in particular to its coupling with streptavidin. In addition, the cellular compartmentalisation of the SNAP tag may allow the development of recombinant markers for inner or outer membranes, which would be a major advantage for the study of outer membrane vesicles, for example.

In summary, we have improved previous methods of markerless gene modification by constructing a cassette combining two counter-selection markers. We have also constructed two new complementation plasmids and demonstrated the feasibility of using hAGT-based technology in *N. meningitidis*. We believe that our tools will be readily applicable to other *Neisseriaceae* and will contribute to the constant need for more tools.

## METHODS

### Strains and growth conditions

*N. meningitidis* 2C4.3 strain is a piliated and capsulated Opa^-^ Opc^-^ variant of the serogroup C meningococcal clinical isolate 8013 (Nassif et al., 1993). *N. meningitidis* Z5463 is a piliated strain isolated from the throat of a patient with meningitis in the Gambia in 1983 (Achtman et al., 1988). All mutants generated in this study are listed in the **supplementary methods**. Bacterial strains were stored frozen at -80°C and routinely grown at 37°C in a moist atmosphere with 5% CO_2_ on gonococcal base medium (GCB) agar plates (Difco) containing Kellogg’s supplements (Kellogg et al., 1968) or in GCB liquid medium [1.5 % proteose peptone (Difco), 0.4 % K_2_HPO_4_, 0.1 % KH_2_PO_4_, 0.1 % NaCl with Kellogg’s supplements]. For selection of meningococcal derivate strains, we used kanamycin at 100 μg/ml, erythromycin (ery) at 2 μg/ml, spectinomycin at 62.5 μg/ml, p-chloro-phenylalanine (Sigma) at 20 mM or 2-deoxy-d-galactose (Sigma) at 1% when it was necessary. Competent strains of *Escherichia coli* DH5-α (New England Biolabs), used for DNA cloning and plasmid propagation, was grown on Lysogeny Broth (LB) agar plates at 37°C or in liquid medium with kanamycin at 50 μg/ml.

### Cell line and growth conditions

EA Hy926 cells (ATCC # CRL-2922) are hybrid endothelial cells, grown in Dulbecco’s modified Eagle’s medium (DMEM) high glucose (Gibco) supplemented with 10% decomplemented fetal calf serum (FCS) at 37°C in a moist atmosphere with 5% CO_2_.

### Transformation of *N. meningitidis*

Strains of interest were grown overnight on GCB agar plates and resuspended to OD_600nm_ = 1 in transformation GCB liquid media containing Kellogg’s supplements, MgCl_2_ at 2.5 mM and MgSO_4_ at 2.5 mM. Five μl of PCR product or 150 ng of plasmid were added to this bacterial suspension and incubated during 30 minutes at 37°C, 5% CO_2_ with agitation, then for 3 hours after addition of 1 ml of transformation GCB liquid media. Transformants were selected on suitable GCB agar plates. Genomic DNA from 12 of the clones was extracted and mutations were verified by PCR using OneTaq polymerase (New England Biolabs) with the appropriate primers (**supplementary methods**) and analyzed by agarose gel electrophoresis.

Genomic DNA was extracted from overnight cultures on GCB agar plates with appropriate components using Wizard Genomic DNA purification kit (Promega) following manufacturer instructions.

### Markerless gene editing in *N. meningitidis*

#### Drop sensitivity assay

Strains of interest grown overnight on GCB agar plates were resuspended to OD_600nm_ = 1 in GCB liquid media containing Kellogg’s supplements. Successive dilutions were deposited on plates containing different concentrations of Cl-Phe and/or 2-DOG.

#### Generation of markerless mutants in *N. meningitidis*

The *aphA(3’)-pheS*-galK* cassette contains the *aphA(3’)* kanamycin resistance gene and the *pheS*\* gene, both under the control of a first promoter and the *galK* gene under the control of a second promoter. This cassette, named APG cassette, was used as follows: **(i)** the first step for the generation of markerless mutant was to introduce the APG selection cassette instead of the ORF (from START codon to STOP codon) of the target gene by homologous recombination. The DNA fragment carrying the cassette was obtained by overlap PCR using Q5 polymerase (New England Biolabs). For each target gene, two DNA fragments ranging from 500 to 1000 base pairs (bp) upstream and downstream of the gene were amplified using two pairs of primers (5’Fw/Rv and 3’Fw/Rv). Another pair of primers (APGFw/Rv) was used to amplify the APG cassette. To enable fragment assembly, the 5’Rv/APGFw primers and the APGRv/3’Fw primers were designed to overlap on 30 nucleotides. Following the amplification of each fragment mentioned above, they were purified using Monarch DNA Gel Extraction kit (New England Biolabs) and assembled by two-step PCR (**supplementary methods**) using Q5 polymerase (New England Biolabs). First, fragments were mixed in stoichiometric quantities and assembled by PCR using “overlap” program: 98°C 30 s, 20 cycles (98°C 20 s, 46°C 1 min, 72°C 1 min/kb), 12°C ∞. Then, the assembled fragment was amplified by PCR with external primers (5’Fw and 3’Rv). Amplified PCR product was introduced into *N. meningitidis* 2C4.3 or Z5463 by transformation and transformants were selected on GCB agar plates containing kanamycin. Transformants sensitivity to Cl-Phe/2- DOG was confirmed on GCB agar + 20 mM Cl-Phe and 1% 2-DOG. Genomic DNA was extracted from 12 transformants, the presence of the APG cassette and the length of fragments bounded by external primers was tested by PCR to confirm the exchange. **(ii)** The second step was to generate markerless mutation. A new DNA fragment constructed by bringing together fragments 5’ and 3’ to delete the entire ORF of the target gene was introduced by homologous recombination instead of the APG cassette. For this, primers 5’RH_Rv and 3’RH_Fw have been modified to overlap on 30 nucleotides. After amplification of each fragment, overlap and amplification of the new fragment as explains above, the fragment was introduced by transformation to a strain of *N. meningitidis* in which the target gene has been replaced by the APG cassette. Transformation is performed as described above, except that the strains of interest are resuspended at OD_600nm_=0.01. Transformants were selected on GCB agar containing 20 mM Cl-Phe and 1% 2-DOG. The absence of the APG cassette was checked by plating on GCB agar + kanamycin and on GCB agar + 20 mM Cl-Phe / 1% 2-DOG and verified by PCR after genomic DNA extraction.

### Liquid growth assays

Strains of interest were grown overnight on GCB agar plates and resuspended to OD_600nm_ = 0.1 in GCB liquid media containing Kellogg’s supplements. Two hundred μl of each bacterial suspension were added into the wells of a 96-well plate and OD_600nm_ was monitored for 24 h at 37°C in 5% CO_2_ using a microplate reader (Tecan Spark). Four independent repeats were carried out for each strain. Curves were obtained using GraphPad Prism (version 8.2) software.

### Construction of pNM99 and pNM42 plasmids

*N. meningitidis* 2C4.3 genomic DNA was used to amplify the DNA fragment comprised between *igA* and *trpB* or *NMV_2142* and *NMV_2143* and the amplicon were inserted in pMR68 (Ramsey et al., 2012) instead of the existing *N. gonorrhoeae* sequence. The resulting plasmids were named pNM42 and pNM99 (primers used are described in the **supplementary methods).**

These plasmids were constructed as follows: the primers were constructed so that they overlapped on 30 bp. The two PCR products were extracted and assembled using the NEBuilder HiFi DNA Assembly kit (New England Biolabs). The assembly product was transformed into *E. coli* DH5-α (New England Biolabs), plated on LB agar plates in the presence of 50 μg/ml kanamycin. The following day, 8 clones were verified by PCR and 4 positive clones were cultured overnight in liquid LB in the presence of 50 μg/ml kanamycin.

Plasmids DNA were extracted (Monarch Plasmid Miniprep, New England Biolabs) and sequenced by Sanger sequencing (Eurofins Genomics). Following the same strategy, these plasmids were then used to insert between *igA*/*trpB* or *NMV_2142*/*NMV_2143* the tetracycline/IPTG-inducible gene expression sequences using pMR33 or pMR68 (Ramsey et al., 2012) as templates and the primers described in **supplementary methods**. The erythromycin resistance gene was exchange with that of spectinomycine only in pNM99 plasmids following the same strategy and with the primers described in **supplementary methods.**

### SNAP-tag expression in *N. meningitidis* 2C4.3

To enable SNAP tag expression in *N. meningitidis*, plasmids pNM99_Ptet_, pNM42_Ptet_, pNM99_Plac_ were used. The plasmids were constructed following the strategy mentioned above. On one side, the SNAP tag (New England Biolabs #N9183S) was inserted after the P*tet* or P*lac* promoter with the appropriate primers (**supplementary methods**). Each ligation product was transformed into *E. coli* DH5-α, extracted and sequenced.

The different plasmids pNM99/42_Ptet/lac_SNAP were introduced into *N. meningitidis* 2C4.3 by transformation. Transformants were selected on GCB agar plates containing erythromycin. The presence of the construct of interest was verified by PCR after genomic DNA extraction.

### Immunoblotting

To detect tag expression by western-blot, one bacterial oese for each strain of interest was resuspended in 500 μl RIPA buffer (50 mM Tris pH 7,5, 150 nM NaCl, 25 nm HEPES, 2 mM EDTA, 1 % wt/vol SDS) after overnight culture in GCB agar + erythromycin + aTC or IPTG at different concentrations to induce expression of the construct. This suspension was heated at 95°C for 5 min. Proteins were quantified using Bradford method and 10 μg of protein was deposited on a 12% acrylamide SDS-PAGE gel. After transfer to nitrocellulose membrane, the membrane was incubated for 20 min in a blocking solution (PBS + tween-20 0.1% + milk 4%) and washed 2 times with PBS + tween-20 0.1%. The membrane was incubated for 1 hour in the presence of an anti-SNAP-tag primary antibody (New England Biolabs #P9310S) diluted at 1:2000 in PBS + tween-20 0.1%, washed 3 times for 5 min with PBS + tween-20 0.1% and then incubated for 45 min with a rabbit anti-IgG secondary antibody coupled to Horse-Raddish Peroxidase (HRP) diluted at 1:10,000 in PBS + tween-20 0.1%. Proteins were detected by chemiluminescence using the Clarity Western ECL substrate detection kit (Bio-Rad) and a Chemidoc imaging system (Bio-rad). Whole proteins were detected on gel using the stain-free technology.

### Immunofluorescence

Four days before infection, 50 000 cells/well were cultured on 2 cm^2^ circular coverslips coated with rat tail type I collagen (Corning). On the day of infection, bacteria grown from an overnight culture on GCB agar plate were adjusted to OD_600nm_=[0.1 and incubated for 2 h at 37[°C in pre-warmed cell culture medium. After 2 hrs incubation at 37°C with 5% CO_2_ under agitation, OD_600nm_ was measured and a total of 10^7^ bacteria were added to cells grown on coverslips and incubated for 40 min at 37°C, 5% CO_2_. After 40 min, the cells were washed 2 times with DMEM and incubated for 2 hrs. Then, cells were washed 2 times with 1X PBS and fixed with 4% paraformaldehyde (PFA) for 20 min, rinsed, incubated 5 min with 50 mM NH_4_Cl and rinsed 2 times with PBS. The coverslips are then incubated for 30 min at 37°C with DMEM supplemented with 10% SVF containing 0.12 μM SNAP-Cell TMR-Star (New England Biolabs #S9105S), rinsed twice with PBS and incubated for 45 min with Dapi (1μg/ml). After 2 additional washings, coverslips were mounted in Mowiol. Image acquisition was performed on Zeiss Apotome fluorescence microscope. Images were collected and processed using the ZEN (ZEISS Efficient Navigation) software.

### Flow cytometry

For quantitative fluorescence analysis, bacteria grown from an overnight culture on GCB agar plates were adjusted to OD_600nm_=0.1 and incubated for 45 min at 37 °C in 5% CO_2_ under agitation in 5 ml pre-warmed GCB liquid media with containing Kellogg’s supplements then incubated with 0.12 μM SNAP-Cell TMR-Star for 30 min at 37°C in 5% CO_2_. A control with no added SNAP-Cell TMR-Star was performed. The suspensions were centrifuged for 5 min at 2,600 g and resuspended in 40 µl of liquid GCB. Ten µl were reintroduced in 3 flasks containing 5 ml of liquid GCB with Kellogg’s supplements and incubated respectively for 30 min, 60 min, and 180 min. The remaining 10 µl were washed twice by centrifugation for 5 min at 1,200 g, then fixed with 500 µl of 4% PFA for 20 min, centrifuged, incubated for 5 min with 50 mM NH_4_Cl, centrifuged, washed 3 times and resuspended in PBS. Recultivated bacteria are respectively centrifuged and fixed after 30 min, 60 min, or 180 min in the same way as above. Two µl of SNAP-Cell Block (New England Biolabs #S9106S) were added to strain 2C4.3 WT. The samples were analysed on FACS (ID700 Spectral Cell Analyzer, Sony) with FlowJo software.

## ACKNOWLEDGEMENTS

This work was supported by the research grant ANR-19-CE14-0045-002 (to MC) and funding from INSERM. Image acquisition and image analysis were performed at the Imaging Facility of Structure Fédérative de Recherche (SFR) Necker, INSERM US24/CNRS UAR3633. Flow cytometry and analysis was performed at the Cytometry facility of SFR Necker, INSERM US24 - CNRS UAR3633. We thank Ana Cehovin and Christoph Tang (Sir William Dunn School of Pathology, University of Oxford, UK) for providing the *galK* cassette.

---

**Supplementary Figure 1.**
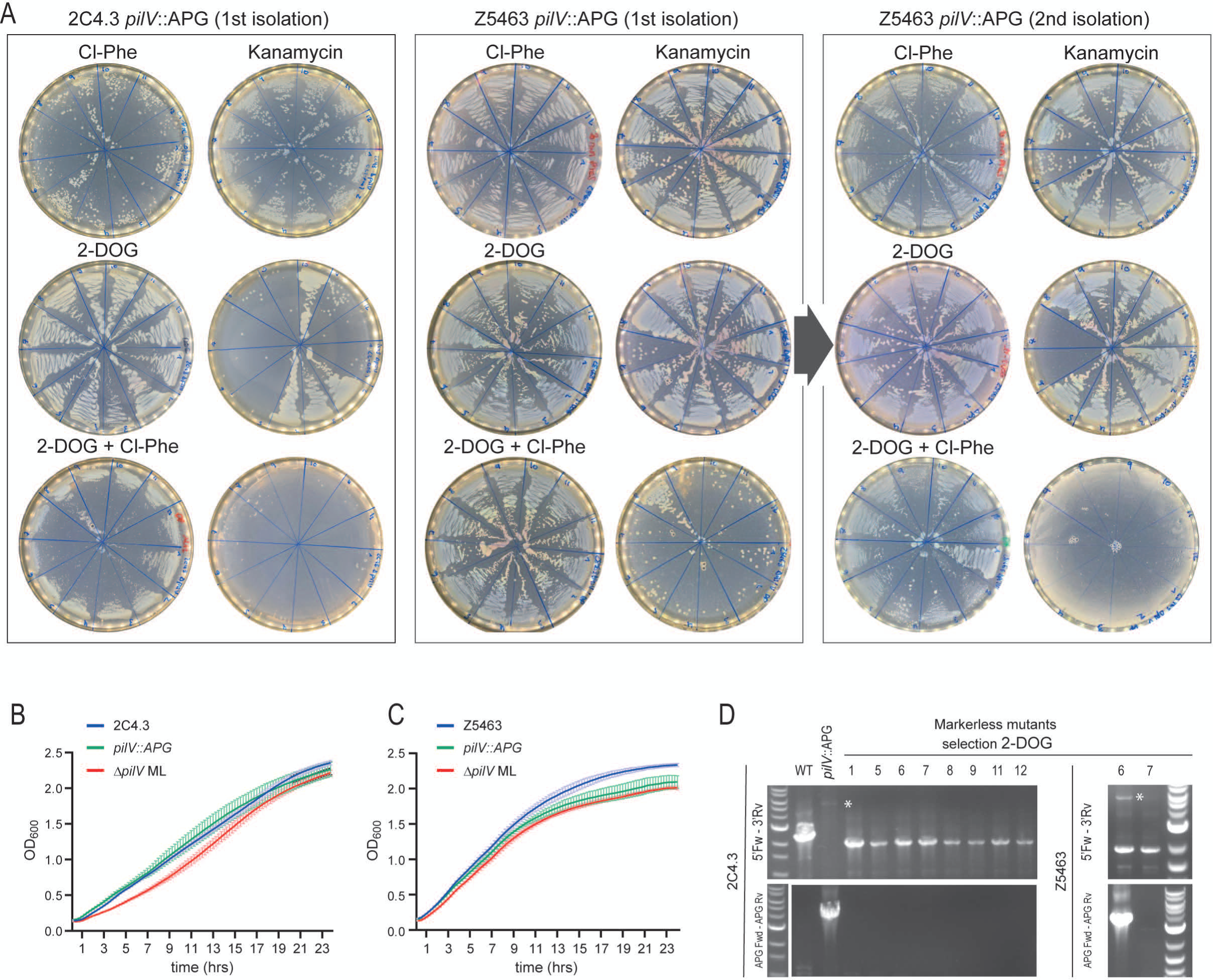
Selection of markerless mutants using 2-deoxy-galactose and p-chloro-phenylalanine or both. **(A)** Selection of markerless mutants on 20 mM Cl-Phe, 1% 2-DOG or both. Mutants carrying the APG cassette in place of the gene *pilV* (*pilV*::APG) were transformed with the fusion product of the 5’ upstream sequence and 3’ downstream sequence of the gene *pilV* to allow homologous recombination and plated on GCB agar plates supplemented with 20 mM Cl-Phe or 1% 2-DOG or both. For each condition and bacterial background, 12 colonies were picked and resuspended and plated again on plates with the same selective compound or with kanamycin. Pictures of plates were then taken after 24 h or 48 h since Cl-Phe may reduce bacterial growth (1^st^ isolation). A second round of selection was carried out for the Z5463 background. For each isolated clone, one colony was picked and isolated again on plates with the same selective compound or with kanamycin (2^nd^ isolation). **(B, C)** Growth of selected mutants is not impaired. The growth of wild type bacteria, a clone of the selected derivative carrying the APG cassette (*pilV*::APG) and a selected markerless clone (Δ*pilV* ML) were assessed in liquid GCB. **(D)** Characterization of markerless mutants in the two backgrounds Z5463 and 2C4.3. WT and derivatives carrying the APG cassette in place of the gene *pilV* (*pilV*::APG) and *pilV* ML mutants obtained by selection on 1% 2-DOG were checked by PCR using the two primer pairs: 5’Fw-3’Rv and APG Fw-APG Rv (Figure 1B). An asterisk indicates the slight band corresponding to the 5’Fw-3’Rv amplicon of the APG. Clone numbers correspond to those on the plates in Supplementary Figure 1A. The largest band on the ladder correspond to 3 kb.

## REFERENCES

1. Achtman, M., Neibert, M., Crowe, B.A., Strittmatter, W., Kusecek, B., Weyse, E., Walsh, M.J., Slawig, B., Morelli, G., Moll, A., et al, 1988. Purification and characterization of eight class 5 outer membrane protein variants from a clone of Neisseria meningitidis serogroup A. J Exp Med 168, 507–25.

2. Alper, M.D., Ames, B.N., 1975. Positive selection of mutants with deletions of the gal-chl region of the Salmonella chromosome as a screening procedure for mutagens that cause deletions. J. Bacteriol. 121, 259–266. 10.1128/jb.121.1.259-266.1975

3. Baerentsen, R., Tang, C.M., Exley, R.M., 2022. Et tu, Neisseria? Conflicts of Interest Between Neisseria Species. Front. Cell. Infect. Microbiol. 12, 913292. 10.3389/fcimb.2022.913292

4. Brynildsrud, O.B., Eldholm, V., Bohlin, J., Uadiale, K., Obaro, S., Caugant, D.A., 2018. Acquisition of virulence genes by a carrier strain gave rise to the ongoing epidemics of meningococcal disease in West Africa. Proc. Natl. Acad. Sci. 115, 5510–5515. 10.1073/pnas.1802298115

5. Cehovin, A., Simpson, P.J., McDowell, M.A., Brown, D.R., Noschese, R., Pallett, M., Brady, J., Baldwin, G.S., Lea, S.M., Matthews, S.J., Pelicic, V., 2013. Specific DNA recognition mediated by a type IV pilin. Proc Natl Acad Sci U A 110, 3065–70. 10.1073/pnas.1218832110

6. Christodoulides, M., Everson, J.S., Liu, B.L., Lambden, P.R., Watt, P.J., Thomas, E.J., Heckels, J.E., 2000. Interaction of primary human endometrial cells with Neisseria gonorrhoeae expressing green fluorescent protein. Mol. Microbiol. 35, 32–43. 10.1046/j.1365-2958.2000.01694.x

7. Facius, D., Fussenegger, M., Meyer, T.F., 1996. Sequential action of factors involved in natural competence for transformation of Neisseria gonorrhoeae. FEMS Microbiol. Lett. 137, 159– 164. 10.1111/j.1574-6968.1996.tb08099.x

8. Gangel, H., Hepp, C., Müller, S., Oldewurtel, E.R., Aas, F.E., Koomey, M., Maier, B., 2014. Concerted spatio-temporal dynamics of imported DNA and ComE DNA uptake protein during gonococcal transformation. PLoS Pathog. 10, e1004043. 10.1371/journal.ppat.1004043

9. Ganini, D., Leinisch, F., Kumar, A., Jiang, J., Tokar, E.J., Malone, C.C., Petrovich, R.M., Mason, R.P., 2017. Fluorescent proteins such as eGFP lead to catalytic oxidative stress in cells. Redox Biol. 12, 462–468. 10.1016/j.redox.2017.03.002

10. Geslewitz, W.E., Cardenas, A., Zhou, X., Zhang, Y., Criss, A.K., Seifert, H.S., 2023. Development and implementation of a Type I-C CRISPR-based programmable repression system for Neisseria gonorrhoeae. mBio e0302523. 10.1128/mbio.03025-23

11. Gurung, I., Berry, J.-L., Hall, A.M.J., Pelicic, V., 2017. Cloning-independent markerless gene editing in Streptococcus sanguinis: novel insights in type IV pilus biology. Nucleic Acids Res. 45, e40. 10.1093/nar/gkw1177

12. Jones, R.A., Yee, W.X., Mader, K., Tang, C.M., Cehovin, A., 2022. Markerless gene editing in Neisseria gonorrhoeae. Microbiol. Read. Engl. 168. 10.1099/mic.0.001201

13. Kast, P., 1994. pKSS--a second-generation general purpose cloning vector for efficient positive selection of recombinant clones. Gene 138, 109–114. 10.1016/0378-1119(94)90790-0

14. Kellogg, D.S., Cohen, I.R., Norins, L.C., Schroeter, A.L., Reising, G., 1968. Neisseria gonorrhoeae. II. Colonial variation and pathogenicity during 35 months in vitro. J Bacteriol 96, 596–605.

15. Keppler, A., Gendreizig, S., Gronemeyer, T., Pick, H., Vogel, H., Johnsson, K., 2003. A general method for the covalent labeling of fusion proteins with small molecules in vivo. Nat. Biotechnol. 21, 86–89. 10.1038/nbt765

16. Laver, J.R., Hughes, S.E., Read, R.C., 2015. Neisserial Molecular Adaptations to the Nasopharyngeal Niche. Adv Microb Physiol 66, 323–55. 10.1016/bs.ampbs.2015.05.001

17. Mairey, E., Genovesio, A., Donnadieu, E., Bernard, C., Jaubert, F., Pinard, E., Seylaz, J., Olivo-Marin, J.C., Nassif, X., Dumenil, G., 2006. Cerebral microcirculation shear stress levels determine Neisseria meningitidis attachment sites along the blood-brain barrier. J Exp Med 203, 1939– 50.

18. Mehr, I.J., Seifert, H.S., 1998. Differential roles of homologous recombination pathways in Neisseria gonorrhoeae pilin antigenic variation, DNA transformation and DNA repair. Mol. Microbiol. 30, 697–710. 10.1046/j.1365-2958.1998.01089.x

19. Nassif, X., Lowy, J., Stenberg, P., O’Gaora, P., Ganji, A., So, M., 1993. Antigenic variation of pilin regulates adhesion of Neisseria meningitidis to human epithelial cells. Mol. Microbiol. 8, 719–725. 10.1111/j.1365-2958.1993.tb01615.x

20. Nyongesa, S., Chenal, M., Bernet, È., Coudray, F., Veyrier, F.J., 2022. Sequential markerless genetic manipulations of species from the Neisseria genus. Can. J. Microbiol. 68, 551–560. 10.1139/cjm-2022-0024

21. Ramsey, M.E., Hackett, K.T., Kotha, C., Dillard, J.P., 2012. New Complementation Constructs for Inducible and Constitutive Gene Expression in Neisseria gonorrhoeae and Neisseria meningitidis. Appl. Environ. Microbiol. 78, 3068–3078. 10.1128/AEM.07871-11

22. Roostalu, J., Jõers, A., Luidalepp, H., Kaldalu, N., Tenson, T., 2008. Cell division in Escherichia coli cultures monitored at single cell resolution. BMC Microbiol. 8, 68. 10.1186/1471-2180-8-68

23. Seow, V.Y., Tsygelnytska, O., Biais, N., 2023. Multisite transformation in Neisseria gonorrhoeae: insights on transformations mechanisms and new genetic modification protocols. Front. Microbiol. 14, 1178128. 10.3389/fmicb.2023.1178128

24. Tobiason, D.M., Seifert, H.S., 2010. Genomic Content of Neisseria Species. J. Bacteriol. 192, 2160– 2168. 10.1128/jb.01593-09

25. Ueki, T., Inouye, S., Inouye, M., 1996. Positive-negative KG cassettes for construction of multi-gene deletions using a single drug marker. Gene 183, 153–157. 10.1016/s0378-1119(96)00546-x

