## supplementary methods for "A Toolbox for *Neisseria meningitidis*: Gene Editing, Complementation and Labelling"

### Primers used

|  | Names | Sequences | Amplification products |
| --- | --- | --- | --- |
| Characterization of the  *aphA(3’)-pheS-galK*  (APG) cassette | PilV_ML_5’_Fw | gccgtctgaaaaacgaaatatcctgccccgc | Region upstream of *pilV* |
|  | PilV_ML_5’_Rv | cgattttgaaaccacagcaatgtgtttccatttgtttgtc |  |
|  | PilV_ML_APG_Fw | tggaaacacattgctgtggtttcaaaatcggctccg | APG resistance cassette |
|  | PilV_ML_APG_Rv | gggttcggcggaggaattcagcactgtcctgctcc |  |
|  | PilV_ML_3’_Fw | aggacagtgctgaattcctccgccgaacccgc | Region downstream of *pilV* |
|  | PilV_ML_3’_Rv | ttcctgaatcactttggcagcg |  |
|  | PilV_RH_ML_5’_Rv | gggttcggcggaggaagcaatgtgtttccatttgtttgtc | Upstream region for HR |
|  | PilV_RH_ML_3’_Fw | tggaaacacattgcttcctccgccgaacccgc | Downstream region for HR |
|  | PilV_Fw | atgaaaaacgttcaaaaaggctttac | *pilV* amplification |
|  | PilV_Rv | tcagtcgaagtccgggcag |  |
| SNAP-tag cloning in pMiniT | PilC1_2C43_Fw | atgaataaaactttaaaaaggcagg | *pilC1* 2C4.3 amplification |
|  | PilC1_2C43_Rv | tcagaagaagacttcacgccagc |  |
|  | 2C43PilC1_tag_Fw | gggctcgaggttaatgcaaataaccccaataagagtacc | Construction pMiniT-(PS-SNAP-tag-C-ter *pilC1* 2C4.3) |
|  | 2C43PilC1_tag_Rv | gcaatctttgtccatgttcatgataatagcgtatttactgg |  |
|  | SNAP_PilC1_tag_Fw | gctattatcatgaacatggacaaagattgcgaaatgaaac |  |
|  | SNAP_PilC1_tag_Rv | attggggttatttgcattaacctcgagcccgggg |  |
| SNAP-tag cloning in pNM plasmids | pNM99_TET_Snap_Fw | ctcgaggttaattaagcggtggcggccgctc | Construction pNM99_Ptet__SNAP-tag |
|  | pNM99_TET_Snap_Rv | gcaatctttgtccatgtggagctccaattggccc |  |
|  | SNAP_Fw | caattggagctccacatggacaaagattgcgaaatgaaac |  |
|  | SNAP_Rv | agcggccgccaccgcttaattaacctcgagcccgggg |  |
|  | pNM99_TET_Snap_Fw | ctcgaggttaattaagcggtggcggccgctc | Construction pNM42_Ptet__SNAP-tag |
|  | PNM42_TET_Rv | gcaatctttgtccatggtggagctccaattggccc |  |
|  | SNAP_TET42_Fw | caattggagctccaccatggacaaagattgcgaaatgaaac |  |
|  | SNAP_Rv | agcggccgccaccgcttaattaacctcgagcccgggg |  |
|  | pNM_LAC_Snap_Fw | ctcgaggttaattaatgaccatgattacgaattcccgg | Construction pNM99_Plac__SNAP-tag |
|  | pNM_LAC_Snap_Rv | gcaatctttgtccattagctgtttcctgtgtgaaattgtt |  |
|  | SNAP_LAC_Fw | cacaggaaacagctaatggacaaagattgcgaaatgaaac |  |
|  | SNAP_LAC_Rv | tcgtaatcatggtcattaattaacctcgagcccgggg |  |
|  | pNM99_TET_NSC_Fw | gaagtcttcttctgagcggtggcggccgctc | Construction pNM99_Ptet__ PS-SNAP-tag-Cter *pilC1* 2C4.3 |
|  | pNM99_TET_NSC_Rv | taaagttttattcatgtggagctccaattggccc |  |
|  | NSC_Fw | caattggagctccacatgaataaaactttaaaaaggcagg |  |
|  | NSC_Rv | agcggccgccaccgctcagaagaagacttcacgccagc |  |
|  | pNM99T_PS_Snap_Rv | attaacctcgagcccggggg | Construction pNM99_Ptet__ PS-SNAP-tag |
|  | pNM99T_PS_Snap_Fw | tgagcggtggcggccgc |  |
| Sequencing | PilC1_2C43_seq1 | gtgcagttgggtgtcttgag | Sequencing *pilC1* 2C4.3 |
|  | PilC1_2C43_seq2 | cgaatccacccttgccaaag |  |
|  | SP6 | atttaggtgacactatag | Sequencing pMiniT |
|  | T7 | taatacgactcactataggg |  |
|  | pNM42/99_tet_seq_Fw | agagtcgaattgttagcggag | Sequencing pNM99_Ptet_ |
|  | pNM99_tet_seq_Rv2 | attttgctatgagtcgaggtcga |  |
|  | pNM42/99_tet_seq_Fw | agagtcgaattgttagcggag | Sequencing pNM42_Ptet_ |
|  | pNM42_tet_seq_Rv | caatatcgaatcggcagggg |  |
|  | M13rev29 | gag cgg ata aca att tca cac agg | Sequencing pNM99_Plac_ |
|  | pNM99_lac_seq_Rv | cctctcaggccgtttaaaca |  |
| Construction of pNM plasmids | pNM_2142_Fw | gatccccaccggaattattatgcccacttccaccaaac | Construction pNM42 |
|  | pNM_2143_Rv | accccgggcggccgccatcccagcgaggacattatgag |  |
|  | pNM_2143pMR68_Fw | gtcctcgctgggatggcggccgcccggggtgggcgaag |  |
|  | pNM_2143_pMR68_Rv | ggaagtgggcataataattccggtggggatcaccg |  |
|  | pNM_igA_Fw | gatccccaccggaattatatcggcgaagaaaacgtgcg | Construction pNM99 |
|  | pNM_igA_Rv | accccgggcggccgcagcttgagaagccggttacaa |  |
|  | pNM_igAK&ori_Fw | ccggcttctcaagctgcggccgcccggggtgggcgaag |  |
|  | pNM_igAK&ori_Rv | tttcttcgccgatataattccggtggggatcaccg |  |
|  | PTET42_Fw | aaacccgtttcggtgtccctttagtgagggttaattgc | Construction pNM42_Ptet_ |
|  | PTET42_Rv | aacacggtttcgataatcgataagcttgatatcgaattcc |  |
|  | pNM42Ptet_Fw | atcaagcttatcgattatcgaaaccgtgttgcccctg |  |
|  | pNM42Ptet_Rv | ccctcactaaagggacaccgaaacgggttttgccga |  |
|  | PTET99_Fw | gggattaaactgtaactttgttagtctgataaaaatgccg | Construction pNM99_Ptet_ |
|  | PTET99_Rv | atcagacggacgaggtccctttagtgagggttaattgc |  |
|  | pNM99Ptet_Fw | ccctcactaaagggacctcgtccgtctgatatagtg |  |
|  | pNM99Ptet_Rv | atcagactaacaaagttacagtttaatccctttgagcttc |  |
|  | PLAC42_Fw | ttgtccgatggggcgctcgaggtcgacggtatcgata | Construction pNM99_Plac_ |
|  | PLAC42_Rv | aacggattccgtttgcgagattttcaggagctatccc |  |
|  | pNM99Plac_Fw | ctcctgaaaatctcgcaaacggaatccgtttatcggc |  |
|  | pNM99Plac_Rv | accgtcgacctcgagcgccccatcggacaaaatgcc |  |
| Exchange of Ery to Spc | pNM99Plac_Spec_Fw | atgaaaaccgccactttataaccctctttattttttcctcc | Construction of pNM99_Plac_/SpecR |
|  | pNM99Plac_Spec_Rv | gtgaggggttttttggttaagggatagctcctgaaaatc |  |
|  | Plac_SpecR_Fw | taaagagggttataaagtggcggttttcatggcttg |  |
|  | Plac_SpecR_Rv | gagctatcccttaaccaaaaaacccctcacgacgc |  |
|  | pNM99Ptet_Spc_Fw | tatgcatcccttaacagtggcggttttcatggcttg | Construction of pNM99_Ptet_/SpecR |
|  | pNM99Ptet_Spc_Rv | taaagagggttataacaaaaaacccctcacgacgc |  |
|  | Ptet_SpecR_Fw | atgaaaaccgccactgttaagggatgcataaactgca |  |
|  | Ptet_SpecR_Rv | ggaggaaaaaataaagagggttataacaaaaaacccctcac |  |

### Strains used

| Strain | Description | Antibiotic resistance | Reference |
| --- | --- | --- | --- |
| 2C4.3 | *N. meningitidis* | / | Nassif *et al*., 1993 |
| Z5463 | *N. meningitidis* | / | Achtman *et al*., 1988 |
| 2C4.3 *pilV*::*apha3’pheSgalk* | *pilV* replaced by APG cassette | Kanamycin | This study |
| 2C4.3 *∆pilV* | Markerless deletion of *pilV* | / | This study |
| Z5463 *pilV*::*apha3’pheSgalk* | *pilV* replaced by APG cassette | Kanamycin | This study |
| Z5463 *∆pilV* | Markerless deletion of *pilV* | / | This study |
| 2C4.3 *99_tet_SNAP-tag* | Surexpression of SNAP-tag by transformation of 2C4.3 with pNM99_Ptet__SNAP-tag | Erythromycin | This study |
| 2C4.3 *42_tet_SNAP-tag* | Surexpression of SNAP-tag by transformation of 2C4.3 with pNM42_Ptet__SNAP-tag | Erythromycin | This study |
| 2C4.3 *99_lac_SNAP-tag* | Surexpression of SNAP-tag by transformation of 2C4.3 with pNM99_Plac__SNAP-tag | Erythromycin | This study |

### Sequence : APG Cassette

LOCUS	Cassette_APG	3529 bp	DNA	linear	18-DEC-2023

DEFINITION .

ACCESSION

VERSION

SOURCE	.

ORGANISM .

COMMENT

COMMENT	ApEinfo:methylated:1

FEATURES	Location/Qualifiers

misc_feature	185..979

/locus_tag="Aph3'"

/label="Aph3'"

/ApEinfo_label="Aph3'"

/ApEinfo_fwdcolor="#ff80ff"

/ApEinfo_revcolor="#ff80ff"

/ApEinfo_graphicformat="arrow_data {{0 0.5 0 1 2 0 0 -1 0

-0.5} {} 0} width 5 offset 0"

misc_feature	990..2048

/locus_tag="pheS"

/label="pheS"

/ApEinfo_label="pheS"

/ApEinfo_fwdcolor="#408080"

/ApEinfo_revcolor="#408080"

/ApEinfo_graphicformat="arrow_data {{0 0.5 0 1 2 0 0 -1 0

-0.5} {} 0} width 5 offset 0"

CDS	2369..3517

/locus_tag="galK"

/label="galK"

/ApEinfo_label="galK"

/ApEinfo_fwdcolor="#339966"

/ApEinfo_revcolor="#339966"

/ApEinfo_graphicformat="arrow_data {{0 0.5 0 1 2 0 0 -1 0

-0.5} {} 0} width 5 offset 0"

misc_feature	2089..2313

/locus_tag="OpaB promoter"

/label="OpaB promoter"

/ApEinfo_label="OpaB promoter"

/ApEinfo_fwdcolor="#ff9900"

/ApEinfo_revcolor="#ff9900"

/ApEinfo_graphicformat="arrow_data {{0 .5 0 -.5} {0 .5 0

-.5} 0} width 5 offset 0"

misc_feature	3520..3529

/locus_tag="DUS"

/label="DUS"

/ApEinfo_label="DUS"

/ApEinfo_fwdcolor="#ff8000"

/ApEinfo_revcolor="#ff8000"

/ApEinfo_graphicformat="arrow_data {{0 0.5 0 1 2 0 0 -1 0

-0.5} {} 0} width 5 offset 0"

misc_feature	complement(1255..1264)

/locus_tag="DUS"

/label="DUS(1)"

/ApEinfo_label="DUS"

/ApEinfo_fwdcolor="#ff8000"

/ApEinfo_revcolor="#ff8000"

/ApEinfo_graphicformat="arrow_data {{0 0.5 0 1 2 0 0 -1 0

-0.5} {} 0} width 5 offset 0"

ORIGIN

1 gtggtttcaa aatcggctcc gtcgatacta tgttatacgc caactttgaa aacaactttg

61 aaaaagctgt tttctggtat ttaaggtttt agaatgcaag gaacagtgaa ttggagttcg

121 tcttgttata attagcttct tggggtatct ttaaatactg tagaaaagag gaaggaaata

181 ataaATGGCT AAAATGAGAA TATCACCGGA ATTGAAAAAA CTGATCGAAA AATACCGCTG

241 CGTAAAAGAT ACGGAAGGAA TGTCTCCTGC TAAGGTATAT AAGCTGGTGG GAGAAAATGA

301 AAACCTATAT TTAAAAATGA CGGACAGCCG GTATAAAGGG ACCACCTATG ATGTGGAACG

361 GGAAAAGGAC ATGATGCTAT GGCTGGAAGG AAAGCTGCCT GTTCCAAAGG TCCTGCACTT

421 TGAACGGCAT GATGGCTGGA GCAATCTGCT CATGAGTGAG GCCGATGGCG TCCTTTGCTC

481 GGAAGAGTAT GAAGATGAAC AAAGCCCTGA AAAGATTATC GAGCTGTATG CGGAGTGCAT

541 CAGGCTCTTT CACTCCATCG ACATATCGGA TTGTCCCTAT ACGAATAGCT TAGACAGCCG

601 CTTAGCCGAA TTGGATTACT TACTGAATAA CGATCTGGCC GATGTGGATT GCGAAAACTG

661 GGAAGAAGAC ACTCCATTTA AAGATCCGCG CGAGCTGTAT GATTTTTTAA AGACGGAAAA

721 GCCCGAAGAG GAACTTGTCT TTTCCCACGG CGACCTGGGA GACAGCAACA TCTTTGTGAA

781 AGATGGCAAA GTAAGTGGCT TTATTGATCT TGGGAGAAGC GGCAGGGCGG ACAAGTGGTA

841 TGACATTGCC TTCTGCGTCC GGTCGATCAG GGAGGATATC GGGGAAGAAC AGTATGTCGA

901 GCTATTTTTT GACTTACTGG GGATCAAGCC TGATTGGGAG AAAATAAAAT ATTATATTTT

961 ACTGGATGAA TTGTTTTAGa ggagtaattA TGTCAGGCTA TACAGGCGGC CTCGTTGTTT

1021 CAGGTGGCAT ATCATTAATT GACAGACTTG ATATTATGGA AAATGTAAAC CGCATCGTTG

1081 CAGAAGGCAT TGCCGCAGTA GAAGCTGCGC AAGACTTCAA CGCTCTAGAA CAAATCAAAG

1141 CCCGTTATCT TGGCAAAACC GGCGAGTTGA CCGGACTTCT GAAAACTTTG GGGCAAATGT

1201 CGCCTGAAGA GCGCAAAACC ATAGGTGCGC ATATCAATGA ATGCAAAAAC CGGTTTCAGA

1261 CGGCTTTTAA TGCCAAACGC GATGCCCTCA ACGAAGTCAA GCTGCAAGCC CGACTTGCCG

1321 CCGAAGCCCT CGATATTACC CTGCCCGGAC GCGCTCAGGA AGGCGGCAGC CTGCATCCCG

1381 TAACCCTGAC CTTGCAACGT GTGGTCGAAC TCTTTCACGG AATGGGTTTC GAAGTGGCGG

1441 ACGGGCCTGA AATCGAAGAC GATTTTCACA ATTTCCAAGC CCTGAACATC CCTGCAAACC

1501 ATCCTGCCCG TGCGATGCAG GATACGTTTT ACGTTGAAAA CGGCGATGTT TTGCGTACGC

1561 ACACTTCCCC GATTCAAATC CGCTATATGC TCGATAAAAA AGAGCCGCCC ATCCGCATTA

1621 TCGCCCCCGG CCGCGTTTAC CGTGTGGACA GCGATGCCAC GCACTCGCCT ATGTTCCATC

1681 AGGCGGAAGG TTTGTGGGTA GAAGAGGGCG TAACTTTTGC CGATTTGAAA GCAGTGTTCA

1741 CGGATTTTAT CCGTCGCTTC TTTGAACGCG ATGATTTGCA AGTACGTTTC CGCCCGTCTT

1801 TCTTCCCGTT CACCGAACCG TCTGCCGAAA TCGACATTAT GGGCGAAAAC GGCAAATGGC

1861 TGGAAGTCGG CGGCTGCGGC ATGGTGCATC CCAACGTGTT GAAAAATGTG AATATCGATT

1921 CTGAAAAATA CACCGGTTTC ggtTTCGGTA TCGGTCTCGA CCGTTTCGCG ATGCTGCGTT

1981 ACAACGTGAA CGATTTGCGC TTGTTTTTTG ATAACGATTT GAATTTTTTG AAGCAGTTTG

2041 TAAAATGAtt gtgtaggctg cccgcctgaa aagaaaatgt gtaccgaggg aatgacggcg

2101 gaaagatgcc cgacggtctt tatagcggat taacaaaaat caggacaagg cggcgggccg

2161 cagacagtac aaatggtacg gaaccgatcc gcccggtgct tcatcacctt agggaaccgt

2221 tccctttgag ccggggcggg gcaacccgta ccggtttttg ttcatccgcc atattgtgtt

2281 gaaacaccgc ccggaacccg atataatccg cccTCATTAA AGAGCAGAAA TTAACTATGA

2341 GAGGATCCAC TATAGCagga gtaattttat gagtctgaaa gaaaaaacac aatctctgtt

2401 tgccaacgca tttggctacc ctgccactca caccattcag gcgcctggcc gcgtgaattt

2461 gattggtgaa cacaccgact acaacgacgg tttcgttctg ccctgcgcga ttgattatca

2521 aaccgtgatc agttgtgcac cacgcgatga ccgtaaagtt cgcgtgatgg cagccgatta

2581 tgaaaatcag ctcgacgagt tttccctcga tgcgcccatt gtcgcacatg aaaactatca

2641 atgggctaac tacgttcgtg gcgtggtgaa acatctgcaa ctgcgtaaca acagcttcgg

2701 cggcgtggac atggtgatca gcggcaatgt gccgcagggt gccgggttaa gttcttccgc

2761 ttcactggaa gtcgcggtcg gaaccgtatt gcagcagctt tatcatctgc cgctggacgg

2821 cgcacaaatc gcgcttaacg gtcaggaagc agaaaaccag tttgtaggct gtaactgcgg

2881 gatcatggat cagctaattt ccgcgctcgg caagaaagat catgccttgc tgatcgattg

2941 ccgctcactg gggaccaaag cagtttccat gcccaaaggt gtggctgtcg tcatcatcaa

3001 cagtaacttc aaacgtaccc tggttggcag cgaatacaac acccgtcgtg aacagtgcga

3061 aaccggtgcg cgtttcttcc agcagccagc cctgcgtgat gtcaccattg aagagttcaa

3121 cgctgttgcg catgaactgg acccgatcgt ggcaaaacgc gtgcgtcata tactgactga

3181 aaacgcccgc accgttgaag ctgccagcgc gctggagcaa ggcgacctga aacgtatggg

3241 cgagttgatg gcggagtctc atgcctctat gcgcgatgat ttcgaaatca ccgtgccgca

3301 aattgacact ctggtagaaa tcgtcaaagc tgtgattggc gacaaaggtg gcgtacgcat

3361 gaccggcggc ggatttggcg gctgtatcgt cgcgctgatc ccggaagagc tggtgcctgc

3421 cgtacagcaa gctgtcgctg aacaatatga agcaaaaaca ggtattaaag agacttttta

3481 cgtttgtaaa ccatcacaag gagcaggaca gtgctgaatg ccgtctgaa

//

### Sequence : pNM42_Ptet_

LOCUS	pNM42_TET	5481 bp	DNA	circular	18-DEC-2023

DEFINITION .

ACCESSION

VERSION

SOURCE	.

ORGANISM .

COMMENT

COMMENT

COMMENT	ApEinfo:methylated:1

FEATURES	Location/Qualifiers

misc_feature	complement(2807..3253)

/locus_tag="2142"

/label="2142"

/ApEinfo_label="2142"

/ApEinfo_fwdcolor="#8080ff"

/ApEinfo_revcolor="#8080ff"

/ApEinfo_graphicformat="arrow_data {{0 1 2 0 0 -1} {} 0}

width 5 offset 0"

misc_feature	4..396

/locus_tag="2143"

/label="2143"

/ApEinfo_label="2143"

/ApEinfo_fwdcolor="#80ff80"

/ApEinfo_revcolor="#80ff80"

/ApEinfo_graphicformat="arrow_data {{0 1 2 0 0 -1} {} 0}

width 5 offset 0"

misc_feature	complement(2739..2798)

/locus_tag="Terminator 42"

/label="Terminator 42"

/ApEinfo_label="Terminator 42"

/ApEinfo_fwdcolor="#0080c0"

/ApEinfo_revcolor="#0080c0"

/ApEinfo_graphicformat="arrow_data {{0 1 2 0 0 -1} {} 0}

width 5 offset 0"

misc_feature	402..434

/locus_tag="terminator 43"

/label="terminator 43"

/ApEinfo_label="terminator 43"

/ApEinfo_fwdcolor="#0080c0"

/ApEinfo_revcolor="#0080c0"

/ApEinfo_graphicformat="arrow_data {{0 1 2 0 0 -1} {} 0}

width 5 offset 0"

misc_feature	887..1366

/locus_tag="ermC"

/label="ermC"

/ApEinfo_label="ermC"

/ApEinfo_fwdcolor="#ffff00"

/ApEinfo_revcolor="#ffff00"

/ApEinfo_graphicformat="arrow_data {{0 1 2 0 0 -1} {} 0}

width 5 offset 0"

misc_feature	633..886

/locus_tag="ermC(1)"

/label="ermC(1)"

/ApEinfo_label="ermC"

/ApEinfo_fwdcolor="#ffff00"

/ApEinfo_revcolor="#ffff00"

/ApEinfo_graphicformat="arrow_data {{0 1 2 0 0 -1} {} 0}

width 5 offset 0"

misc_feature	2487..2607

/locus_tag="Promoter tet"

/label="Promoter tet"

/ApEinfo_label="Promoter tet"

/ApEinfo_fwdcolor="#008080"

/ApEinfo_revcolor="#008080"

/ApEinfo_graphicformat="arrow_data {{0 1 2 0 0 -1} {} 0}

width 5 offset 0"

misc_feature	complement(2226..2291)

/locus_tag="TetR"

/label="TetR"

/ApEinfo_label="TetR"

/ApEinfo_fwdcolor="#008080"

/ApEinfo_revcolor="#008080"

/ApEinfo_graphicformat="arrow_data {{0 1 2 0 0 -1} {} 0}

width 5 offset 0"

misc_feature	complement(1668..2225)

/locus_tag="TetR(1)"

/label="TetR(1)"

/ApEinfo_label="TetR"

/ApEinfo_fwdcolor="#008080"

/ApEinfo_revcolor="#008080"

/ApEinfo_graphicformat="arrow_data {{0 1 2 0 0 -1} {} 0}

width 5 offset 0"

CDS	complement(3455..4246)

/locus_tag="Kan/neoR"

/label="Kan/neoR"

/ApEinfo_label="Kan/neoR"

/ApEinfo_fwdcolor="yellow"

/ApEinfo_revcolor="yellow"

/ApEinfo_graphicformat="arrow_data {{0 1 2 0 0 -1} {} 0}

width 5 offset 0"

misc_feature	403..412

/locus_tag="DUS"

/label="DUS"

/ApEinfo_label="DUS"

/ApEinfo_fwdcolor="#ff8080"

/ApEinfo_revcolor="#ff8080"

/ApEinfo_graphicformat="arrow_data {{0 1 2 0 0 -1} {} 0}

width 5 offset 0"

misc_feature	1442..1451

/locus_tag="DUS(1)"

/label="DUS(1)"

/ApEinfo_label="DUS"

/ApEinfo_fwdcolor="#ff8080"

/ApEinfo_revcolor="#ff8080"

/ApEinfo_graphicformat="arrow_data {{0 1 2 0 0 -1} {} 0}

width 5 offset 0"

misc_feature	2751..2760

/locus_tag="DUS(2)"

/label="DUS(2)"

/ApEinfo_label="DUS"

/ApEinfo_fwdcolor="#ff8080"

/ApEinfo_revcolor="#ff8080"

/ApEinfo_graphicformat="arrow_data {{0 1 2 0 0 -1} {} 0}

width 5 offset 0"

misc_feature	complement(424..433)

/locus_tag="DUS(3)"

/label="DUS(3)"

/ApEinfo_label="DUS"

/ApEinfo_fwdcolor="#ff8080"

/ApEinfo_revcolor="#ff8080"

/ApEinfo_graphicformat="arrow_data {{0 1 2 0 0 -1} {} 0}

width 5 offset 0"

misc_feature	complement(1457..1466)

/locus_tag="DUS(4)"

/label="DUS(4)"

/ApEinfo_label="DUS"

/ApEinfo_fwdcolor="#ff8080"

/ApEinfo_revcolor="#ff8080"

/ApEinfo_graphicformat="arrow_data {{0 1 2 0 0 -1} {} 0}

width 5 offset 0"

misc_feature	complement(2775..2784)

/locus_tag="DUS(5)"

/label="DUS(5)"

/ApEinfo_label="DUS"

/ApEinfo_fwdcolor="#ff8080"

/ApEinfo_revcolor="#ff8080"

/ApEinfo_graphicformat="arrow_data {{0 1 2 0 0 -1} {} 0}

width 5 offset 0"

misc_feature	4247..4280

/locus_tag="NeoKan promoter"

/label="NeoKan promoter"

/ApEinfo_label="NeoKan promoter"

/ApEinfo_fwdcolor="#80ff80"

/ApEinfo_revcolor="#80ff80"

/ApEinfo_graphicformat="arrow_data {{0 1 2 0 0 -1} {} 0}

width 5 offset 0"

misc_feature	4831..5449

/locus_tag="ORI"

/label="ORI"

/ApEinfo_label="ORI"

/ApEinfo_fwdcolor="#ffa87d"

/ApEinfo_revcolor="#ffa87d"

/ApEinfo_graphicformat="arrow_data {{0 1 2 0 0 -1} {} 0}

width 5 offset 0"

misc_feature	2633..2633

/locus_tag="Best insertion site"

/label="Best insertion site"

/ApEinfo_label="Best insertion site"

/ApEinfo_fwdcolor="#ff0000"

/ApEinfo_revcolor="#ff0000"

/ApEinfo_graphicformat="arrow_data {{0 1 2 0 0 -1} {} 0}

width 5 offset 0"

ORIGIN

1 attatgccca cttccaccaa accctacatc ctccgcgccc tctgcgaatg gtgcagcgac

61 aacagcctca caccgcacat ccttgtctgg gtcaacgaac acacgcgcgt ccccatgcag

121 tacgtccgcg acaacgaaat tatgctcaac atcggcgcga ccgccacgca aaaccttcaa

181 atcgacaacg attggatcag tttttccgcc cgtttcagcg gacaggcgca cgatatatgg

241 atacctgtcg gacacgtcct cagccttttc gcacgggaga ccggcgaagg tatggggttt

301 gaattggaag cgtatcgccc cgacacgccg cctgaaaaca cctctgccga aaccgcgccc

361 cggtccgcca aaaaaggctt gaaattggtc aaataaatct atgccgtctg aacggaatcg

421 tgtttcagac ggcattttgt ccgatggggc gcaaacggaa tccgtttatc ggcaaaaccc

481 gtttcggtgT CCCTTTAGTG AGGGTTAATT GCGCGCTTGG CGTAATCATG GTCATAGCTG

541 TTCGATAAGC TAATTCTCAT GTTTGACAGC TTATCATCGC GTGCTATAAT TATACTAATT

601 TTATAAGGAG GAAAAAATAA AGAGGGTTAT AATGAACGAG AAAAATATAA AACACAGTCA

661 AAACTTTATT ACTTCAAAAC ATAATATAGA TAAAATAATG ACAAATATAA GATTAAATGA

721 ACATGATAAT ATCTTTGAAA TCGGCTCAGG AAAAGGGCAT TTTACCCTTG AATTAGTACA

781 GAGGTGTAAT TTCGTAACTG CCATTGAAAT AGACCATAAA TTATGCAAAA CTACAGAAAA

841 TAAACTTGTT GATCACGATA ATTTCCAAGT TTTAAACAAG GATATATTGC AGTTTAAATT

901 TCCTAAAAAC CAATCCTATA AAATATTTGG TAATATACCT TATAACATAA GTACGGATAT

961 AATACGCAAA ATTGTTTTTG ATAGTATAGC TGATGAGATT TATTTAATCG TGGAATACGG

1021 GTTTGCTAAA AGATTATTAA ATACAAAACG CTCATTGGCA TTATTTTTAA TGGCAGAAGT

1081 TGATATTTCT ATATTAAGTA TGGTTCCAAG AGAATATTTT CATCCTAAAC CTAAAGTGAA

1141 TAGCTCACTT ATCAGATTAA ATAGAAAAAA ATCAAGAATA TCACACAAAG ATAAACAGAA

1201 GTATAATTAT TTCGTTATGA AATGGGTTAA CAAAGAATAC AAGAAAATAT TTACAAAAAA

1261 TCAATTTAAC AATTCCTTAA AACATGCAGG AATTGACGAT TTAAACAATA TTAGCTTTGA

1321 ACAATTCTTA TCTCTTTTCA ATAGCTATAA ATTATTTAAT AAGTAAGTTA AGGGATGCAT

1381 AAACTGCATC CCTTAACTTG TTTTTCGTGT ACCTATTTTT TGTGAATCGA CTCTAGAGCT

1441 TGCCGTCTGA AATGGTTTCA GACGGCATGC AGCAATGGCA ACAACGTTGA AGCTAGCTGC

1501 GATGAGTGGC AGGGCGGGGC GTAATTTTTT TAAGGCAGTT ATTGGTGCCC TTAAACGCCT

1561 GGTTGCTACG CCTGAATAAG TGATAATAAG CGGATGAATG GCAGAAATTC GAACTCGACA

1621 TCTTGGTTAC CGTGAAGTTA CCATCACGGA AAAAGGTTAT GCTGCTTTTA AGACCCACTT

1681 TCACATTTAA GTTGTTTTTC TAATCCGCAT ATGATCAATT CAAGGCCGAA TAAGAAGGCT

1741 GGCTCTGCAC CTTGGTGATC AAATAATTCG ATAGCTTGTC GTAATAATGG CGGCATACTA

1801 TCAGTAGTAG GTGTTTCCCT TTCTTCTTTA GCGACTTGAT GCTCTTGATC TTCCAATACG

1861 CAACCTAAAG TAAAATGCCC CACAGCGCTG AGTGCATATA ATGCATTCTC TAGTGAAAAA

1921 CCTTGTTGGC ATAAAAAGGC TAATTGATTT TCGAGAGTTT CATACTGTTT TTCTGTAGGC

1981 CGTGTACCTA AATGTACTTT TGCTCCATCG CGATGACTTA GTAAAGCACA TCTAAAACTT

2041 TTAGCGTTAT TACGTAAAAA ATCTTGCCAG CTTTCCCCTT CTAAAGGGCA AAAGTGAGTA

2101 TGGTGCCTAT CTAACATCTC AATGGCTAAG GCGTCGAGCA AAGCCCGCTT ATTTTTTACA

2161 TGCCAATACA ATGTAGGCTG CTCTACACCT AGCTTCTGGG CGAGTTTACG GGTTGTTAAA

2221 CCTTCGATTC CGACCTCATT AAGCAGCTCT AATGCGCTGT TAATCACTTT ACTTTTATCT

2281 AATCTAGACA TTTGATATGC CTCCTAAATT TTTATCTAAA GTGAATTTAG GAGGCTTACT

2341 TGTCTGCTTT CTTCATTAGA ATCAATCCTT TTTTAAAAGT CAATATTACT GTAACATAAA

2401 TATATATTTT AAAAATATCC CACTTTATCC AATTTTCGTT TGTTGAACCA TTATATCACA

2461 TTATCCATTA AAAATCAAAC AAATTTGGAT CTTAAAAAAT TTATTTGACA CCCTATCAGT

2521 GATAGAGTAT AATAGAGTCG AATTGTTAGC GGAGAAGAAT TTCACACAGA ATTCATTAAA

2581 GAGGAGAAAT TAACTATGAG AGGATCCACT ATAGGGCCAA TTGGAGCTCC ACCGCGGTGG

2641 CGGCCGCTCT AGAACTAGTG GATCCCCCGG GCTGCAGGAA TTCGATATCA AGCTTATCGA

2701 Ttatcgaaac cgtgttgccc ctgccgattc gatattggga cattgaaaat gccgtctgaa

2761 cctgcgatac gggcttcaga cggcattttg tccgatattc gggcaatcag gcggtcagta

2821 cggctttcag aattttgttg acttcgccca tatcggcttt gccggcgagg cgggttttca

2881 atacgcccat cactttgccc atatccgcca tacctgccgc gccggtttcg gcaacggcgg

2941 catcgaccgc agtacggatt tcgccggcgg agagcatttg cggcaggtag cggtgcagta

3001 cctcgatttc ggcgttttct ttgtctgcca aatcctgacg gccggcttca gtgtagattt

3061 tcgcgctgtc tttgcgctgt ttgaccattt tggtcaggat ggcggtgatt ttggcatcgt

3121 cggcttcggt gcgttcgtcc acttcaaact gtttgacggc ggcgttgatg aggcggatgg

3181 tgccgaggga aacttggtct ttggcgcgca tcgcggtttt catgtcttcg gtaaggcgga

3241 ttttcaggct cataatgtcc tcgctgggat gGCGGCCGCC CGGGGTGGGC GAAGAACTCC

3301 AGCATGAGAT CCCCGCGCTG GAGGATCATC CAGCCGGCGT CCCGGAAAAC GATTCCGAAG

3361 CCCAACCTTT CATAGAAGGC GGCGGTGGAA TCGAAATCTC GTGATGGCAG GTTGGGCGTC

3421 GCTTGGTCGG TCATTTCGAA CCCCAGAGTC CCGCTCAGAA GAACTCGTCA AGAAGGCGAT

3481 AGAAGGCGAT GCGCTGCGAA TCGGGAGCGG CGATACCGTA AAGCACGAGG AAGCGGTCAG

3541 CCCATTCGCC GCCAAGCTCT TCAGCAATAT CACGGGTAGC CAACGCTATG TCCTGATAGC

3601 GGTCCGCCAC ACCCAGCCGG CCACAGTCGA TGAATCCAGA AAAGCGGCCA TTTTCCACCA

3661 TGATATTCGG CAAGCAGGCA TCGCCATGGG TCACGACGAG ATCCTCGCCG TCGGGCATGC

3721 GCGCCTTGAG CCTGGCGAAC AGTTCGGCTG GCGCGAGCCC CTGATGCTCT TCGTCCAGAT

3781 CATCCTGATC GACAAGACCG GCTTCCATCC GAGTACGTGC TCGCTCGATG CGATGTTTCG

3841 CTTGGTGGTC GAATGGGCAG GTAGCCGGAT CAAGCGTATG CAGCCGCCGC ATTGCATCAG

3901 CCATGATGGA TACTTTCTCG GCAGGAGCAA GGTGAGATGA CAGGAGATCC TGCCCCGGCA

3961 CTTCGCCCAA TAGCAGCCAG TCCCTTCCCG CTTCAGTGAC AACGTCGAGC ACAGCTGCGC

4021 AAGGAACGCC CGTCGTGGCC AGCCACGATA GCCGCGCTGC CTCGTCCTGC AGTTCATTCA

4081 GGGCACCGGA CAGGTCGGTC TTGACAAAAA GAACCGGGCG CCCCTGCGCT GACAGCCGGA

4141 ACACGGCGGC ATCAGAGCAG CCGATTGTCT GTTGTGCCCA GTCATAGCCG AATAGCCTCT

4201 CCACCCAAGC GGCCGGAGAA CCTGCGTGCA ATCCATCTTG TTCAATCATG CGAAACGATC

4261 CTCATCCTGT CTCTTGATCA GATCTTGATC CCCTGCGCCA TCAGATCCTT GGCGGCAAGA

4321 AAGCCATCCA GTTTACTTTG CAGGGCTTCC CAACCTTACC AGAGGGCGCC CCAGCTGGCA

4381 ATTCCGGTTC GCTTGCTGTC CATAAAACCG CCCAGTCTAG CTATCGCCAT GTAAGCCCAC

4441 TGCAAGCTAC CTGCTTTCTC TTTGCGCTTG CGTTTTCCCT TGTCCAGATA GCCCAGTAGC

4501 TGACATTCAT CCGGGGTCAG CACCGTTTCT GCGGACTGGC TTTCTACGTG TTCCGCTTCC

4561 TTTAGCAGCC CTTGCGCCCT GAGTGCTTGC GGCAGCGTGA AGCTAGCTTA TGCGGTGTGA

4621 AATACCGCAC AGATGCGTAA GGAGAAAATA CCGCATCAGG CGCTCTTCCG CTTCCTCGCT

4681 CACTGACTCG CTGCGCTCGG TCGTTCGGCT GCGGCGAGCG GTATCAGCTC ACTCAAAGGC

4741 GGTAATACGG TTATCCACAG AATCAGGGGA TAACGCAGGA AAGAACATGT GAGCAAAAGG

4801 CCAGCAAAAG GCCAGGAACC GTAAAAAGGC CGCGTTGCTG GCGTTTTTCC ATAGGCTCCG

4861 CCCCCCTGAC GAGCATCACA AAAATCGACG CTCAAGTCAG AGGTGGCGAA ACCCGACAGG

4921 ACTATAAAGA TACCAGGCGT TTCCCCCTGG AAGCTCCCTC GTGCGCTCTC CTGTTCCGAC

4981 CCTGCCGCTT ACCGGATACC TGTCCGCCTT TCTCCCTTCG GGAAGCGTGG CGCTTTCTCA

5041 TAGCTCACGC TGTAGGTATC TCAGTTCGGT GTAGGTCGTT CGCTCCAAGC TGGGCTGTGT

5101 GCACGAACCC CCCGTTCAGC CCGACCGCTG CGCCTTATCC GGTAACTATC GTCTTGAGTC

5161 CAACCCGGTA AGACACGACT TATCGCCACT GGCAGCAGCC ACTGGTAACA GGATTAGCAG

5221 AGCGAGGTAT GTAGGCGGTG CTACAGAGTT CTTGAAGTGG TGGCCTAACT ACGGCTACAC

5281 TAGAAGGACA GTATTTGGTA TCTGCGCTCT GCTGAAGCCA GTTACCTTCG GAAAAAGAGT

5341 TGGTAGCTCT TGATCCGGCA AACAAACCAC CGCTGGTAGC GGCGGTTTTT TGTTTGCAAG

5401 CAGCAGATTA CGCGCAGAAA AAAAGGATCT CAAGAAGATC CTTTGATCTT TTCTTACTGA

5461 ACGGTGATCC CCACCGGAAT T

//

### Sequence : pNM99_Ptet_

LOCUS	pNM99_TET	5617 bp	DNA	circular	18-DEC-2023

DEFINITION .

ACCESSION

VERSION

SOURCE	.

ORGANISM .

COMMENT

COMMENT	ApEinfo:methylated:1

FEATURES	Location/Qualifiers

misc_feature	2211..2690

/locus_tag="IgA"

/label="IgA"

/ApEinfo_label="IgA"

/ApEinfo_fwdcolor="#8080ff"

/ApEinfo_revcolor="#8080ff"

/ApEinfo_graphicformat="arrow_data {{0 1 2 0 0 -1} {} 0}

width 5 offset 0"

misc_feature	complement(5160..5617)

/locus_tag="TrpB"

/label="TrpB"

/ApEinfo_label="TrpB"

/ApEinfo_fwdcolor="#8080ff"

/ApEinfo_revcolor="#8080ff"

/ApEinfo_graphicformat="arrow_data {{0 1 2 0 0 -1} {} 0}

width 5 offset 0"

misc_feature	complement(2705..2754)

/locus_tag="Terminator"

/label="Terminator"

/ApEinfo_label="Terminator"

/ApEinfo_fwdcolor="#ff80ff"

/ApEinfo_revcolor="#ff80ff"

/ApEinfo_graphicformat="arrow_data {{0 1 2 0 0 -1} {} 0}

width 5 offset 0"

misc_feature	3196..3744

/locus_tag="ermC"

/label="ermC"

/ApEinfo_label="ermC"

/ApEinfo_fwdcolor="#ffff00"

/ApEinfo_revcolor="#ffff00"

/ApEinfo_graphicformat="arrow_data {{0 1 2 0 0 -1} {} 0}

width 5 offset 0"

misc_feature	4865..4985

/locus_tag="Promoter tet"

/label="Promoter tet"

/ApEinfo_label="Promoter tet"

/ApEinfo_fwdcolor="#008080"

/ApEinfo_revcolor="#008080"

/ApEinfo_graphicformat="arrow_data {{0 1 2 0 0 -1} {} 0}

width 5 offset 0"

misc_feature	5107..5143

/locus_tag="terminator"

/label="terminator"

/ApEinfo_label="terminator"

/ApEinfo_fwdcolor="#0000ff"

/ApEinfo_revcolor="#0000ff"

/ApEinfo_graphicformat="arrow_data {{0 1 2 0 0 -1} {} 0}

width 5 offset 0"

misc_feature	3011..3195

/locus_tag="ermC(1)"

/label="ermC(1)"

/ApEinfo_label="ermC"

/ApEinfo_fwdcolor="#ffff00"

/ApEinfo_revcolor="#ffff00"

/ApEinfo_graphicformat="arrow_data {{0 1 2 0 0 -1} {} 0}

width 5 offset 0"

misc_feature	complement(4046..4669)

/locus_tag="TetR"

/label="TetR"

/ApEinfo_label="TetR"

/ApEinfo_fwdcolor="#008080"

/ApEinfo_revcolor="#008080"

/ApEinfo_graphicformat="arrow_data {{0 1 2 0 0 -1} {} 0}

width 5 offset 0"

CDS	1236..2027

/locus_tag="Kan/neoR"

/label="Kan/neoR"

/ApEinfo_label="Kan/neoR"

/ApEinfo_fwdcolor="yellow"

/ApEinfo_revcolor="yellow"

/ApEinfo_graphicformat="arrow_data {{0 1 2 0 0 -1} {} 0}

width 5 offset 0"

misc_feature	complement(3835..3844)

/locus_tag="DUS"

/label="DUS"

/ApEinfo_label="DUS"

/ApEinfo_fwdcolor="#ff8080"

/ApEinfo_revcolor="#ff8080"

/ApEinfo_graphicformat="arrow_data {{0 1 2 0 0 -1} {} 0}

width 5 offset 0"

misc_feature	complement(2735..2744)

/locus_tag="DUS(1)"

/label="DUS(1)"

/ApEinfo_label="DUS"

/ApEinfo_fwdcolor="#ff8080"

/ApEinfo_revcolor="#ff8080"

/ApEinfo_graphicformat="arrow_data {{0 1 2 0 0 -1} {} 0}

width 5 offset 0"

misc_feature	5112..5121

/locus_tag="DUS(2)"

/label="DUS(2)"

/ApEinfo_label="DUS"

/ApEinfo_fwdcolor="#ff8080"

/ApEinfo_revcolor="#ff8080"

/ApEinfo_graphicformat="arrow_data {{0 1 2 0 0 -1} {} 0}

width 5 offset 0"

misc_feature	3820..3829

/locus_tag="DUS(3)"

/label="DUS(3)"

/ApEinfo_label="DUS"

/ApEinfo_fwdcolor="#ff8080"

/ApEinfo_revcolor="#ff8080"

/ApEinfo_graphicformat="arrow_data {{0 1 2 0 0 -1} {} 0}

width 5 offset 0"

misc_feature	2718..2727

/locus_tag="DUS(4)"

/label="DUS(4)"

/ApEinfo_label="DUS"

/ApEinfo_fwdcolor="#ff8080"

/ApEinfo_revcolor="#ff8080"

/ApEinfo_graphicformat="arrow_data {{0 1 2 0 0 -1} {} 0}

width 5 offset 0"

misc_feature	complement(1202..1235)

/locus_tag="NeoKan promoter"

/label="NeoKan promoter"

/ApEinfo_label="NeoKan promoter"

/ApEinfo_fwdcolor="#80ff80"

/ApEinfo_revcolor="#80ff80"

/ApEinfo_graphicformat="arrow_data {{0 1 2 0 0 -1} {} 0}

width 5 offset 0"

misc_feature	complement(33..651)

/locus_tag="ORI"

/label="ORI"

/ApEinfo_label="ORI"

/ApEinfo_fwdcolor="#ffa87d"

/ApEinfo_revcolor="#ffa87d"

/ApEinfo_graphicformat="arrow_data {{0 1 2 0 0 -1} {} 0}

width 5 offset 0"

misc_feature	complement(5011..5011)

/locus_tag="Best insertion site"

/label="Best insertion site"

/ApEinfo_label="Best insertion site"

/ApEinfo_fwdcolor="#c0c0c0"

/ApEinfo_revcolor="cyan"

/ApEinfo_graphicformat="arrow_data {{0 1 2 0 0 -1} {} 0}

width 5 offset 0"

ORIGIN

1 AATTCCGGTG GGGATCACCG TTCAGTAAGA AAAGATCAAA GGATCTTCTT GAGATCCTTT

61 TTTTCTGCGC GTAATCTGCT GCTTGCAAAC AAAAAACCGC CGCTACCAGC GGTGGTTTGT

121 TTGCCGGATC AAGAGCTACC AACTCTTTTT CCGAAGGTAA CTGGCTTCAG CAGAGCGCAG

181 ATACCAAATA CTGTCCTTCT AGTGTAGCCG TAGTTAGGCC ACCACTTCAA GAACTCTGTA

241 GCACCGCCTA CATACCTCGC TCTGCTAATC CTGTTACCAG TGGCTGCTGC CAGTGGCGAT

301 AAGTCGTGTC TTACCGGGTT GGACTCAAGA CGATAGTTAC CGGATAAGGC GCAGCGGTCG

361 GGCTGAACGG GGGGTTCGTG CACACAGCCC AGCTTGGAGC GAACGACCTA CACCGAACTG

421 AGATACCTAC AGCGTGAGCT ATGAGAAAGC GCCACGCTTC CCGAAGGGAG AAAGGCGGAC

481 AGGTATCCGG TAAGCGGCAG GGTCGGAACA GGAGAGCGCA CGAGGGAGCT TCCAGGGGGA

541 AACGCCTGGT ATCTTTATAG TCCTGTCGGG TTTCGCCACC TCTGACTTGA GCGTCGATTT

601 TTGTGATGCT CGTCAGGGGG GCGGAGCCTA TGGAAAAACG CCAGCAACGC GGCCTTTTTA

661 CGGTTCCTGG CCTTTTGCTG GCCTTTTGCT CACATGTTCT TTCCTGCGTT ATCCCCTGAT

721 TCTGTGGATA ACCGTATTAC CGCCTTTGAG TGAGCTGATA CCGCTCGCCG CAGCCGAACG

781 ACCGAGCGCA GCGAGTCAGT GAGCGAGGAA GCGGAAGAGC GCCTGATGCG GTATTTTCTC

841 CTTACGCATC TGTGCGGTAT TTCACACCGC ATAAGCTAGC TTCACGCTGC CGCAAGCACT

901 CAGGGCGCAA GGGCTGCTAA AGGAAGCGGA ACACGTAGAA AGCCAGTCCG CAGAAACGGT

961 GCTGACCCCG GATGAATGTC AGCTACTGGG CTATCTGGAC AAGGGAAAAC GCAAGCGCAA

1021 AGAGAAAGCA GGTAGCTTGC AGTGGGCTTA CATGGCGATA GCTAGACTGG GCGGTTTTAT

1081 GGACAGCAAG CGAACCGGAA TTGCCAGCTG GGGCGCCCTC TGGTAAGGTT GGGAAGCCCT

1141 GCAAAGTAAA CTGGATGGCT TTCTTGCCGC CAAGGATCTG ATGGCGCAGG GGATCAAGAT

1201 CTGATCAAGA GACAGGATGA GGATCGTTTC GCATGATTGA ACAAGATGGA TTGCACGCAG

1261 GTTCTCCGGC CGCTTGGGTG GAGAGGCTAT TCGGCTATGA CTGGGCACAA CAGACAATCG

1321 GCTGCTCTGA TGCCGCCGTG TTCCGGCTGT CAGCGCAGGG GCGCCCGGTT CTTTTTGTCA

1381 AGACCGACCT GTCCGGTGCC CTGAATGAAC TGCAGGACGA GGCAGCGCGG CTATCGTGGC

1441 TGGCCACGAC GGGCGTTCCT TGCGCAGCTG TGCTCGACGT TGTCACTGAA GCGGGAAGGG

1501 ACTGGCTGCT ATTGGGCGAA GTGCCGGGGC AGGATCTCCT GTCATCTCAC CTTGCTCCTG

1561 CCGAGAAAGT ATCCATCATG GCTGATGCAA TGCGGCGGCT GCATACGCTT GATCCGGCTA

1621 CCTGCCCATT CGACCACCAA GCGAAACATC GCATCGAGCG AGCACGTACT CGGATGGAAG

1681 CCGGTCTTGT CGATCAGGAT GATCTGGACG AAGAGCATCA GGGGCTCGCG CCAGCCGAAC

1741 TGTTCGCCAG GCTCAAGGCG CGCATGCCCG ACGGCGAGGA TCTCGTCGTG ACCCATGGCG

1801 ATGCCTGCTT GCCGAATATC ATGGTGGAAA ATGGCCGCTT TTCTGGATTC ATCGACTGTG

1861 GCCGGCTGGG TGTGGCGGAC CGCTATCAGG ACATAGCGTT GGCTACCCGT GATATTGCTG

1921 AAGAGCTTGG CGGCGAATGG GCTGACCGCT TCCTCGTGCT TTACGGTATC GCCGCTCCCG

1981 ATTCGCAGCG CATCGCCTTC TATCGCCTTC TTGACGAGTT CTTCTGAGCG GGACTCTGGG

2041 GTTCGAAATG ACCGACCAAG CGACGCCCAA CCTGCCATCA CGAGATTTCG ATTCCACCGC

2101 CGCCTTCTAT GAAAGGTTGG GCTTCGGAAT CGTTTTCCGG GACGCCGGCT GGATGATCCT

2161 CCAGCGCGGG GATCTCATGC TGGAGTTCTT CGCCCACCCC GGGCGGCCGC agcttgagaa

2221 gccggttaca aacgcagcaa aaagcaaact ttaaccgaac aagcatccaa accggcctta

2281 ctttgggcaa tacgctgaaa atcaatcaat tcgagattgt ccccagtgcg ggaatccgtt

2341 acagccgcct gtcatctgca gattacaagt tgggtaacga cagtgttaaa gtaagttcta

2401 tgtcagtgaa aacactgacg gccggactgg attttgctta tcggtttaaa gtcggcaacc

2461 ttaccgtaaa acccttgtta tctgcagctt actttgccaa ttatggcaaa ggcggcgtga

2521 atgtgggcgg taattccttc gcctataaag cagataatca acagcaatat tcagcaggtg

2581 ccgcgttact gtaccgtaat gttacattaa acgtaaatgg cagtattaca aaaggcaaac

2641 aattggaaaa acaaaaatcc ggacaaatta aaatacagat tcgtttctaa aatactaaat

2701 tcatagcaaa ataaaatgcc gtctgaactc aggcttcaga cggcattttt atagtggatt

2761 aacaaaaatc aggacaaggc gacgaagccG CAGACAGTAC AGATAGACGG CAAGGCGAGG

2821 CAACGCCGTA CTGGTTTTTG TTAATCcact atatcagacg gacgaggTCC CTTTAGTGAG

2881 GGTTAATTGC GCGCTTGGCG TAATCATGGT CATAGCTGTT CGATAAGCTA ATTCTCATGT

2941 TTGACAGCTT ATCATCGCGT GCTATAATTA TACTAATTTT ATAAGGAGGA AAAAATAAAG

3001 AGGGTTATAA TGAACGAGAA AAATATAAAA CACAGTCAAA ACTTTATTAC TTCAAAACAT

3061 AATATAGATA AAATAATGAC AAATATAAGA TTAAATGAAC ATGATAATAT CTTTGAAATC

3121 GGCTCAGGAA AAGGGCATTT TACCCTTGAA TTAGTACAGA GGTGTAATTT CGTAACTGCC

3181 ATTGAAATAG ACCATAAATT ATGCAAAACT ACAGAAAATA AACTTGTTGA TCACGATAAT

3241 TTCCAAGTTT TAAACAAGGA TATATTGCAG TTTAAATTTC CTAAAAACCA ATCCTATAAA

3301 ATATTTGGTA ATATACCTTA TAACATAAGT ACGGATATAA TACGCAAAAT TGTTTTTGAT

3361 AGTATAGCTG ATGAGATTTA TTTAATCGTG GAATACGGGT TTGCTAAAAG ATTATTAAAT

3421 ACAAAACGCT CATTGGCATT ATTTTTAATG GCAGAAGTTG ATATTTCTAT ATTAAGTATG

3481 GTTCCAAGAG AATATTTTCA TCCTAAACCT AAAGTGAATA GCTCACTTAT CAGATTAAAT

3541 AGAAAAAAAT CAAGAATATC ACACAAAGAT AAACAGAAGT ATAATTATTT CGTTATGAAA

3601 TGGGTTAACA AAGAATACAA GAAAATATTT ACAAAAAATC AATTTAACAA TTCCTTAAAA

3661 CATGCAGGAA TTGACGATTT AAACAATATT AGCTTTGAAC AATTCTTATC TCTTTTCAAT

3721 AGCTATAAAT TATTTAATAA GTAAGTTAAG GGATGCATAA ACTGCATCCC TTAACTTGTT

3781 TTTCGTGTAC CTATTTTTTG TGAATCGACT CTAGAGCTTG CCGTCTGAAA TGGTTTCAGA

3841 CGGCATGCAG CAATGGCAAC AACGTTGAAG CTAGCTGCGA TGAGTGGCAG GGCGGGGCGT

3901 AATTTTTTTA AGGCAGTTAT TGGTGCCCTT AAACGCCTGG TTGCTACGCC TGAATAAGTG

3961 ATAATAAGCG GATGAATGGC AGAAATTCGA ACTCGACATC TTGGTTACCG TGAAGTTACC

4021 ATCACGGAAA AAGGTTATGC TGCTTTTAAG ACCCACTTTC ACATTTAAGT TGTTTTTCTA

4081 ATCCGCATAT GATCAATTCA AGGCCGAATA AGAAGGCTGG CTCTGCACCT TGGTGATCAA

4141 ATAATTCGAT AGCTTGTCGT AATAATGGCG GCATACTATC AGTAGTAGGT GTTTCCCTTT

4201 CTTCTTTAGC GACTTGATGC TCTTGATCTT CCAATACGCA ACCTAAAGTA AAATGCCCCA

4261 CAGCGCTGAG TGCATATAAT GCATTCTCTA GTGAAAAACC TTGTTGGCAT AAAAAGGCTA

4321 ATTGATTTTC GAGAGTTTCA TACTGTTTTT CTGTAGGCCG TGTACCTAAA TGTACTTTTG

4381 CTCCATCGCG ATGACTTAGT AAAGCACATC TAAAACTTTT AGCGTTATTA CGTAAAAAAT

4441 CTTGCCAGCT TTCCCCTTCT AAAGGGCAAA AGTGAGTATG GTGCCTATCT AACATCTCAA

4501 TGGCTAAGGC GTCGAGCAAA GCCCGCTTAT TTTTTACATG CCAATACAAT GTAGGCTGCT

4561 CTACACCTAG CTTCTGGGCG AGTTTACGGG TTGTTAAACC TTCGATTCCG ACCTCATTAA

4621 GCAGCTCTAA TGCGCTGTTA ATCACTTTAC TTTTATCTAA TCTAGACATT TGATATGCCT

4681 CCTAAATTTT TATCTAAAGT GAATTTAGGA GGCTTACTTG TCTGCTTTCT TCATTAGAAT

4741 CAATCCTTTT TTAAAAGTCA ATATTACTGT AACATAAATA TATATTTTAA AAATATCCCA

4801 CTTTATCCAA TTTTCGTTTG TTGAACCATT ATATCACATT ATCCATTAAA AATCAAACAA

4861 ATTTGGATCT TAAAAAATTT ATTTGACACC CTATCAGTGA TAGAGTATAA TAGAGTCGAA

4921 TTGTTAGCGG AGAAGAATTT CACACAGAAT TCATTAAAGA GGAGAAATTA ACTATGAGAG

4981 GATCCACTAT AGGGCCAATT GGAGCTCCAC CGCGGTGGCG GCCGCTCTAG AACTAGTGGA

5041 TCCCCCGGGC TGCAGGAATT CGATATCAAG CTTATCGATA CCGTCGACCT CGACTCATAG

5101 CAAAATAAAA TGCCGTCTGA ACTCAAGCTT CGGACGGCAT TTTTATCAGA CTAACAAAGt

5161 tacagtttaa tccctttgag cttcgcgacg gtattgatgt ctttgtcgcc acgaccggat

5221 aggttgacca gaatcacttg gtctttgccc attttcggcg cgtttttcac cgcccaagca

5281 acggcgtggc tggattccag cgcaggaata atgccctcga agcggcagag caagtcaaag

5341 gcttcgagtg cttcgtcgtc tttggcaaca gtgtattcga cgcgcttgat gtcgtgcaga

5401 tggctgtgtt ccgggccgat gccggggtaa tccaagcctg cggaaacaga gtgcgtaccc

5461 aaaacctgac cgttttcgtc ctgcatcaga tagctgcgga aaccgtgcaa tacgccaatc

5521 ggtgcgcccg aagtaatcgg cgcggcgtga tcgggggtgt tcacgcccaa accgccagcc

5581 tccacgccga cgaggcgcac gttttcttcg ccgatat

//

### Sequence : pNM99_Plac_

LOCUS	pNM99_LAC	5942 bp	DNA	circular	18-DEC-2023

DEFINITION .

ACCESSION

VERSION

SOURCE	.

ORGANISM .

COMMENT

COMMENT	ApEinfo:methylated:1

FEATURES	Location/Qualifiers

misc_feature	complement(3253..3732)

/locus_tag="IgA"

/label="IgA"

/ApEinfo_label="IgA"

/ApEinfo_fwdcolor="#8080ff"

/ApEinfo_revcolor="#8080ff"

/ApEinfo_graphicformat="arrow_data {{0 1 2 0 0 -1} {} 0}

width 5 offset 0"

misc_feature	1..458

/locus_tag="TrpB"

/label="TrpB"

/ApEinfo_label="TrpB"

/ApEinfo_fwdcolor="#8080ff"

/ApEinfo_revcolor="#8080ff"

/ApEinfo_graphicformat="arrow_data {{0 1 2 0 0 -1} {} 0}

width 5 offset 0"

misc_feature	complement(475..511)

/locus_tag="terminator trpB"

/label="terminator trpB"

/ApEinfo_label="terminator trpB"

/ApEinfo_fwdcolor="#0080ff"

/ApEinfo_revcolor="#0080ff"

/ApEinfo_graphicformat="arrow_data {{0 1 2 0 0 -1} {} 0}

width 5 offset 0"

misc_feature	complement(3191..3242)

/locus_tag="terminator"

/label="terminator"

/ApEinfo_label="terminator"

/ApEinfo_fwdcolor="#0080ff"

/ApEinfo_revcolor="#0080ff"

/ApEinfo_graphicformat="arrow_data {{0 1 2 0 0 -1} {} 0}

width 5 offset 0"

misc_feature	complement(1553..2635)

/locus_tag="LacI"

/label="LacI"

/ApEinfo_label="LacI"

/ApEinfo_fwdcolor="#ff8040"

/ApEinfo_revcolor="#ff8000"

/ApEinfo_graphicformat="arrow_data {{0 1 2 0 0 -1} {} 0}

width 5 offset 0"

misc_feature	complement(2636..2713)

/locus_tag="LacIQ Promoter"

/label="LacIQ Promoter"

/ApEinfo_label="LacIQ Promoter"

/ApEinfo_fwdcolor="#0000a0"

/ApEinfo_revcolor="#0000a0"

/ApEinfo_graphicformat="arrow_data {{0 1 2 0 0 -1} {} 0}

width 5 offset 0"

misc_feature	complement(769..1375)

/locus_tag="ErmC"

/label="ErmC"

/ApEinfo_label="ErmC"

/ApEinfo_fwdcolor="#ffff00"

/ApEinfo_revcolor="green"

/ApEinfo_graphicformat="arrow_data {{0 1 2 0 0 -1} {} 0}

width 5 offset 0"

misc_feature	complement(642..768)

/locus_tag="ErmC(1)"

/label="ErmC(1)"

/ApEinfo_label="ErmC"

/ApEinfo_fwdcolor="#ffff00"

/ApEinfo_revcolor="green"

/ApEinfo_graphicformat="arrow_data {{0 1 2 0 0 -1} {} 0}

width 5 offset 0"

misc_feature	4708..4741

/locus_tag="NeoKAN promoter"

/label="NeoKAN promoter"

/ApEinfo_label="NeoKAN promoter"

/ApEinfo_fwdcolor="#80ff80"

/ApEinfo_revcolor="#80ff80"

/ApEinfo_graphicformat="arrow_data {{0 1 2 0 0 -1} {} 0}

width 5 offset 0"

misc_binding	2975..2997

/locus_tag="LacO"

/label="LacO"

/ApEinfo_label="LacO"

/ApEinfo_fwdcolor="#6495ed"

/ApEinfo_revcolor="#6495ed"

/ApEinfo_graphicformat="arrow_data {{0 1 2 0 0 -1} {} 0}

width 5 offset 0"

CDS	complement(3916..4707)

/locus_tag="Kan/neoR"

/label="Kan/neoR"

/ApEinfo_label="Kan/neoR"

/ApEinfo_fwdcolor="yellow"

/ApEinfo_revcolor="yellow"

/ApEinfo_graphicformat="arrow_data {{0 1 2 0 0 -1} {} 0}

width 5 offset 0"

misc_feature	1462..1471

/locus_tag="DUS"

/label="DUS"

/ApEinfo_label="DUS"

/ApEinfo_fwdcolor="#ff8080"

/ApEinfo_revcolor="#ff8080"

/ApEinfo_graphicformat="arrow_data {{0 1 2 0 0 -1} {} 0}

width 5 offset 0"

misc_feature	complement(497..506)

/locus_tag="DUS(1)"

/label="DUS(1)"

/ApEinfo_label="DUS"

/ApEinfo_fwdcolor="#ff8080"

/ApEinfo_revcolor="#ff8080"

/ApEinfo_graphicformat="arrow_data {{0 1 2 0 0 -1} {} 0}

width 5 offset 0"

misc_feature	4708..4741

/locus_tag="NeoKan promoter"

/label="NeoKan promoter"

/ApEinfo_label="NeoKan promoter"

/ApEinfo_fwdcolor="#80ff80"

/ApEinfo_revcolor="#80ff80"

/ApEinfo_graphicformat="arrow_data {{0 1 2 0 0 -1} {} 0}

width 5 offset 0"

misc_feature	5292..5910

/locus_tag="ORI"

/label="ORI"

/ApEinfo_label="ORI"

/ApEinfo_fwdcolor="#ffa87d"

/ApEinfo_revcolor="#ffa87d"

/ApEinfo_graphicformat="arrow_data {{0 1 2 0 0 -1} {} 0}

width 5 offset 0"

misc_feature	3015..3015

/locus_tag="Best insertion site"

/label="Best insertion site"

/ApEinfo_label="Best insertion site"

/ApEinfo_fwdcolor="#ff0000"

/ApEinfo_revcolor="#ff0000"

/ApEinfo_graphicformat="arrow_data {{0 1 2 0 0 -1} {} 0}

width 5 offset 0"

ORIGIN

1 atatcggcga agaaaacgtg cgcctcgtcg gcgtggaggc tggcggtttg ggcgtgaaca

61 cccccgatca cgccgcgccg attacttcgg gcgcaccgat tggcgtattg cacggtttcc

121 gcagctatct gatgcaggac gaaaacggtc aggttttggg tacgcactct gtttccgcag

181 gcttggatta ccccggcatc ggcccggaac acagccatct gcacgacatc aagcgcgtcg

241 aatacactgt tgccaaagac gacgaagcac tcgaagcctt tgacttgctc tgccgcttcg

301 agggcattat tcctgcgctg gaatccagcc acgccgttgc ttgggcggtg aaaaacgcgc

361 cgaaaatggg caaagaccaa gtgattctgg tcaacctatc cggtcgtggc gacaaagaca

421 tcaataccgt cgcgaagctc aaagggatta aactgtaaCT TTGTTAGTCT GATAAAAATG

481 CCGTCCGAAG CTTGAGTTCA GACGGCATTT TATTTTGCTA TGAGGTACCT TGGTCATGGC

541 CAGCTTATCG GTAAGAGGTT CCAACTTTCA CCATAATGAA ATAAAATCAC TACCGGGCGT

601 ATTTTTTGAG TTATCGAGAT TTTCAGGAGC TATCCCTTAA CTTACTTATT AAATAATTTA

661 TAGCTATTGA AAAGAGATAA GAATTGTTCA AAGCTAATAT TGTTTAAATC GTCAATTCCT

721 GCATGTTTTA AGGAATTGTT AAATTGATTT TTTGTAAATA TTTTCTTGTA TTCTTTGTTA

781 ACCCATTTCA TAACGAAATA ATTATACTTC TGTTTATCTT TGTGTGATAT TCTTGATTTT

841 TTTCTATTTA ATCTGATAAG TGAGCTATTC ACTTTAGGTT TAGGATGAAA ATATTCTCTT

901 GGAACCATAC TTAATATAGA AATATCAACT TCTGCCATTA AAAATAATGC CAATGAGCGT

961 TTTGTATTTA ATAATCTTTT AGCAAACCCG TATTCCACGA TTAAATAAAT CTCATCAGCT

1021 ATACTATCAA AAACAATTTT GCGTATTATA TCCGTACTTA TGTTATAAGG TATATTACCA

1081 AATATTTTAT AGGATTGGTT TTTAGGAAAT TTAAACTGCA ATATATCCTT GTTTAAAACT

1141 TGGAAATTAT CGTGATCAAC AAGTTTATTT TCTGTAGTTT TGCATAATTT ATGGTCTATT

1201 TCAATGGCAG TTACGAAATT ACACCTCTGT ACTAATTCAA GGGTAAAATG CCCTTTTCCT

1261 GAGCCGATTT CAAAGATATT ATCATGTTCA TTTAATCTTA TATTTGTCAT TATTTTATCT

1321 ATATTATGTT TTGAAGTAAT AAAGTTTTGA CTGTGTTTTA TATTTTTCTC GTTCATTATA

1381 ACCCTCTTTA TTTTTTCCTC CTTATAAAAT TAGTATAATT ATAGCACGCG ATGATAAGCT

1441 GTCAAACATG AGAATTAGCT TGCCGTCTGA AATGGCCCGA ACGCCAGCAA GACGTAGCCC

1501 AGCGCGTCGG CCAGCTTGCA ATTCGCGCTA ACTTACATTA ATTGCGTTGC GCTCACTGCC

1561 CGCTTTCCAG TCGGGAAACC TGTCGTGCCA GCTGCATTAA TGAATCGGCC AACGCGCGGG

1621 GAGAGGCGGT TTGCGTATTG GGCGCCAGGG TGGTTTTTCT TTTCACCAGT GAGACGGGCA

1681 ACAGCTGATT GCCCTTCACC GCCTGGCCCT GAGAGAGTTG CAGCAAGCGG TCCACGCTGG

1741 TTTGCCCCAG CAGGCGAAAA TCCTGTTTGA TGGTGGTTAA CGGCGGGATA TAACATGAGC

1801 TGTCTTCGGT ATCGTCGTAT CCCACTACCG AGATATCCGC ACCAACGCGC AGCCCGGACT

1861 CGGTAATGGC GCGCATTGCG CCCAGCGCCA TCTGATCGTT GGCAACCAGC ATCGCAGTGG

1921 GAACGATGCC CTCATTCAGC ATTTGCATGG TTTGTTGAAA ACCGGACATG GCACTCCAGT

1981 CGCCTTCCCG TTCCGCTATC GGCTGAATTT GATTGCGAGT GAGATATTTA TGCCAGCCAG

2041 CCAGACGCAG ACGCGCCGAG ACAGAACTTA ATGGGCCCGC TAACAGCGCG ATTTGCTGGT

2101 GACCCAATGC GACCAGATGC TCCACGCCCA GTCGCGTACC GTCTTCATGG GAGAAAATAA

2161 TACTGTTGAT GGGTGTCTGG TCAGAGACAT CAAGAAATAA CGCCGGAACA TTAGTGCAGG

2221 CAGCTTCCAC AGCAATGGCA TCCTGGTCAT CCAGCGGATA GTTAATGATC AGCCCACTGA

2281 CGCGTTGCGC GAGAAGATTG TGCACCGCCG CTTTACAGGC TTCGACGCCG CTTCGTTCTA

2341 CCATCGACAC CACCACGCTG GCACCCAGTT GATCGGCGCG AGATTTAATC GCCGCGACAA

2401 TTTGCGACGG CGCGTGCAGG GCCAGACTGG AGGTGGCAAC GCCAATCAGC AACGACTGTT

2461 TGCCCGCCAG TTGTTGTGCC ACGCGGTTGG GAATGTAATT CAGCTCCGCC ATCGCCGCTT

2521 CCACTTTTTC CCGCGTTTTC GCAGAAACGT GGCTGGCCTG GTTCACCACG CGGGAAACGG

2581 TCTGATAAGA GACACCGGCA TACTCTGCGA CATCGTATAA CGTTACTGGT TTCACATTCA

2641 CCACCCTGAA TTGACTCTCT TCCGGGCGCT ATCATGCCAT ACCGCGAAAG GTTTTGCACC

2701 ATTCGATGGT GTCAACGTAA ATGCCGCTTC GCCTTCGCGC GCGAATTGCA AGCTGATCCG

2761 GGCTTATCGA CTGCACGGTG CACCAATGCT TCTGGCGTCA GGCAGCCATC GGAAGCTGTG

2821 GTATGGCTGT GCAGGTCGTA AATCACTGCA TAATTCGTGT CGCTCAAGGC GCACTCCCGT

2881 TCTGGATAAT GTTTTTTGCG CCGACATCAT AACGGTTCTG GCAAATATTC TGAAATGAGC

2941 TGTTGACAAT TAATCATCGG CTCGTATAAT GTGTGGAATT GTGAGCGGAT AACAATTTCA

3001 CACAGGAAAC AGCTATGACC ATGATTACGA ATTCCCGGAT TAATTAAGTA CTATGCATGT

3061 TTAAACGGCC TGAGAGGATC CACTATAGGG CCAATTGGAG CTCCACCGCG GTGGCGGCCG

3121 CTCTAGAACT AGTGGATCCC CCGGGCTGCA GGAATTCGAT ATCAAGCTTA TCGATACCGT

3181 CGACCTCGAG GGGGGGCCCG GTACCCCCTT AAGGGATGCA GTTTATGCAT CCCTTAAATT

3241 TAGTATTgta ttttagaaac gaatctgtat tttaatttgt ccggattttt gtttttccaa

3301 ttgtttgcct tttgtaatac tgccatttac gtttaatgta acattacggt acagtaacgc

3361 ggcacctgct gaatattgct gttgattatc tgctttatag gcgaaggaat taccgcccac

3421 attcacgccg cctttgccat aattggcaaa gtaagctgca gataacaagg gttttacggt

3481 aaggttgccg actttaaacc gataagcaaa atccagtccg gccgtcagtg ttttcactga

3541 catagaactt actttaacac tgtcgttacc caacttgtaa tctgcagatg acaggcggct

3601 gtaacggatt cccgcactgg ggacaatctc gaattgattg attttcagcg tattgcccaa

3661 agtaaggccg gtttggatgc ttgttcggtt aaagtttgct ttttgctgcg tttgtaaccg

3721 gcttctcaag ctGCGGCCGC CCGGGGTGGG CGAAGAACTC CAGCATGAGA TCCCCGCGCT

3781 GGAGGATCAT CCAGCCGGCG TCCCGGAAAA CGATTCCGAA GCCCAACCTT TCATAGAAGG

3841 CGGCGGTGGA ATCGAAATCT CGTGATGGCA GGTTGGGCGT CGCTTGGTCG GTCATTTCGA

3901 ACCCCAGAGT CCCGCTCAGA AGAACTCGTC AAGAAGGCGA TAGAAGGCGA TGCGCTGCGA

3961 ATCGGGAGCG GCGATACCGT AAAGCACGAG GAAGCGGTCA GCCCATTCGC CGCCAAGCTC

4021 TTCAGCAATA TCACGGGTAG CCAACGCTAT GTCCTGATAG CGGTCCGCCA CACCCAGCCG

4081 GCCACAGTCG ATGAATCCAG AAAAGCGGCC ATTTTCCACC ATGATATTCG GCAAGCAGGC

4141 ATCGCCATGG GTCACGACGA GATCCTCGCC GTCGGGCATG CGCGCCTTGA GCCTGGCGAA

4201 CAGTTCGGCT GGCGCGAGCC CCTGATGCTC TTCGTCCAGA TCATCCTGAT CGACAAGACC

4261 GGCTTCCATC CGAGTACGTG CTCGCTCGAT GCGATGTTTC GCTTGGTGGT CGAATGGGCA

4321 GGTAGCCGGA TCAAGCGTAT GCAGCCGCCG CATTGCATCA GCCATGATGG ATACTTTCTC

4381 GGCAGGAGCA AGGTGAGATG ACAGGAGATC CTGCCCCGGC ACTTCGCCCA ATAGCAGCCA

4441 GTCCCTTCCC GCTTCAGTGA CAACGTCGAG CACAGCTGCG CAAGGAACGC CCGTCGTGGC

4501 CAGCCACGAT AGCCGCGCTG CCTCGTCCTG CAGTTCATTC AGGGCACCGG ACAGGTCGGT

4561 CTTGACAAAA AGAACCGGGC GCCCCTGCGC TGACAGCCGG AACACGGCGG CATCAGAGCA

4621 GCCGATTGTC TGTTGTGCCC AGTCATAGCC GAATAGCCTC TCCACCCAAG CGGCCGGAGA

4681 ACCTGCGTGC AATCCATCTT GTTCAATCAT GCGAAACGAT CCTCATCCTG TCTCTTGATC

4741 AGATCTTGAT CCCCTGCGCC ATCAGATCCT TGGCGGCAAG AAAGCCATCC AGTTTACTTT

4801 GCAGGGCTTC CCAACCTTAC CAGAGGGCGC CCCAGCTGGC AATTCCGGTT CGCTTGCTGT

4861 CCATAAAACC GCCCAGTCTA GCTATCGCCA TGTAAGCCCA CTGCAAGCTA CCTGCTTTCT

4921 CTTTGCGCTT GCGTTTTCCC TTGTCCAGAT AGCCCAGTAG CTGACATTCA TCCGGGGTCA

4981 GCACCGTTTC TGCGGACTGG CTTTCTACGT GTTCCGCTTC CTTTAGCAGC CCTTGCGCCC

5041 TGAGTGCTTG CGGCAGCGTG AAGCTAGCTT ATGCGGTGTG AAATACCGCA CAGATGCGTA

5101 AGGAGAAAAT ACCGCATCAG GCGCTCTTCC GCTTCCTCGC TCACTGACTC GCTGCGCTCG

5161 GTCGTTCGGC TGCGGCGAGC GGTATCAGCT CACTCAAAGG CGGTAATACG GTTATCCACA

5221 GAATCAGGGG ATAACGCAGG AAAGAACATG TGAGCAAAAG GCCAGCAAAA GGCCAGGAAC

5281 CGTAAAAAGG CCGCGTTGCT GGCGTTTTTC CATAGGCTCC GCCCCCCTGA CGAGCATCAC

5341 AAAAATCGAC GCTCAAGTCA GAGGTGGCGA AACCCGACAG GACTATAAAG ATACCAGGCG

5401 TTTCCCCCTG GAAGCTCCCT CGTGCGCTCT CCTGTTCCGA CCCTGCCGCT TACCGGATAC

5461 CTGTCCGCCT TTCTCCCTTC GGGAAGCGTG GCGCTTTCTC ATAGCTCACG CTGTAGGTAT

5521 CTCAGTTCGG TGTAGGTCGT TCGCTCCAAG CTGGGCTGTG TGCACGAACC CCCCGTTCAG

5581 CCCGACCGCT GCGCCTTATC CGGTAACTAT CGTCTTGAGT CCAACCCGGT AAGACACGAC

5641 TTATCGCCAC TGGCAGCAGC CACTGGTAAC AGGATTAGCA GAGCGAGGTA TGTAGGCGGT

5701 GCTACAGAGT TCTTGAAGTG GTGGCCTAAC TACGGCTACA CTAGAAGGAC AGTATTTGGT

5761 ATCTGCGCTC TGCTGAAGCC AGTTACCTTC GGAAAAAGAG TTGGTAGCTC TTGATCCGGC

5821 AAACAAACCA CCGCTGGTAG CGGCGGTTTT TTGTTTGCAA GCAGCAGATT ACGCGCAGAA

5881 AAAAAGGATC TCAAGAAGAT CCTTTGATCT TTTCTTACTG AACGGTGATC CCCACCGGAA

5941 TT

//

### Sequence : aadA Cassette

LOCUS	cassette_aadA	1070 bp	DNA	linear	18-DEC-2023

DEFINITION .

ACCESSION

VERSION

SOURCE	.

ORGANISM .

COMMENT

COMMENT

COMMENT	ApEinfo:methylated:1

FEATURES	Location/Qualifiers

misc_feature	164..952

/locus_tag="aadA"

/label="aadA"

/ApEinfo_label="aadA"

/ApEinfo_fwdcolor="#ffb83e"

/ApEinfo_revcolor="#2fffff"

/ApEinfo_graphicformat="arrow_data {{0 1 2 0 0 -1} {} 0}

width 5 offset 0"

misc_feature	13..62

/locus_tag="promoter"

/label="promoter"

/ApEinfo_label="promoter"

/ApEinfo_fwdcolor="#ffc919"

/ApEinfo_revcolor="#5dff4f"

/ApEinfo_graphicformat="arrow_data {{0 1 2 0 0 -1} {} 0}

width 5 offset 0"

ORIGIN

1 AGTGGCGGTT TTCATGGCTT GTTATGACTG TTTTTTTGGG GTACAGTCTA TGCCTCGGGC

61 ATCCAAGCAG CAAGCGCGTT ACGCCGTGGG TCGATGTTTG ATGTTATGGA GCAGCAACGA

121 TGTTACGCAG CAGGGCAGTC GCCCTAAAAC AAAGTTAAAC ATCATGAGGG AAGCGGTGAT

181 CGCCGAAGTA TCGACTCAAC TATCAGAGGT AGTTGGCGTC ATCGAGCGCC ATCTCGAACC

241 GACGTTGCTG GCCGTACATT TGTACGGCTC CGCAGTGGAT GGCGGCCTGA AGCCACACAG

301 TGATATTGAT TTGCTGGTTA CGGTGACCGT AAGGCTTGAT GAAACAACGC GGCGAGCTTT

361 GATCAACGAC CTTTTGGAAA CTTCGGCTTC CCCTGGAGAG AGCGAGATTC TCCGCGCTGT

421 AGAAGTCACC ATTGTTGTGC ACGACGACAT CATTCCGTGG CGTTATCCAG CTAAGCGCGA

481 ACTGCAATTT GGAGAATGGC AGCGCAATGA CATTCTTGCA GGTATCTTCG AGCCAGCCAC

541 GATCGACATT GATCTGGCTA TCTTGCTGAC AAAAGCAAGA GAACATAGCG TTGCCTTGGT

601 AGGTCCAGCG GCGGAGGAAC TCTTTGATCC GGTTCCTGAA CAGGATCTAT TTGAGGCGCT

661 AAATGAAACC TTAACGCTAT GGAACTCGCC GCCCGACTGG GCTGGCGATG AGCGAAATGT

721 AGTGCTTACG TTGTCCCGCA TTTGGTACAG CGCAGTAACC GGCAAAATCG CGCCGAAGGA

781 TGTCGCTGCC GACTGGGCAA TGGAGCGCCT GCCGGCCCAG TATCAGCCCG TCATACTTGA

841 AGCTAGGCAG GCTTATCTTG GACAAGAAGA TCGCTTGGCC TCGCGCGCAG ATCAGTTGGA

901 AGAATTTGTT CACTACGTGA AAGGCGAGAT CACCAAGGTA GTCGGCAAAT AATGTCTAAC

961 AATTCGTTCA AGCCGACGCC GCTTCGCGGC GCGGCTTAAC TCAAGCGTAG CAATCACTAA

1021 GTAGGATAAC CCCCTGGGGG CCTCTCCCCG GCGTCGTGAG GGGTTTTTTG

//
